# Cetuximab increases LGR5 expression and augments LGR5-targeting antibody-drug conjugate efficacy in patient-derived colorectal cancer models

**DOI:** 10.1101/2025.06.18.660406

**Authors:** Peyton C. High, Zhengdong Liang, Cara Guernsey-Biddle, Shraddha Subramanian, Yueh-Ming Shyu, Adela M. Aldana, Yukimatsu Toh, Kendra S. Carmon

**Affiliations:** Center for Translational Cancer Research, The Brown Foundation Institute of Molecular Medicine, University of Texas Health Science Center at Houston, Houston, TX, 77030, USA; The University of Texas MD Anderson Cancer Center UTHealth Houston Graduate School of Biomedical Sciences, Houston, TX, 77030, USA

**Keywords:** Antibody-drug conjugate, LGR5, EGFR, colorectal cancer, combination treatment, cetuximab

## Abstract

Colorectal cancer (CRC) remains the second-leading cause of cancer-associated deaths, indicating an urgent need for improved therapeutic options. We previously generated antibody-drug conjugates (ADCs) targeting the cancer stem-like cell marker leucine-rich repeat-containing G protein-coupled receptor 5 (LGR5). However, tumor relapse due to LGR5 downregulation and suboptimal payload selection warranted strategies to improve ADC efficacy. Here we report cetuximab, an EGFR-targeting monoclonal antibody indicated for RAS^WT^ metastatic CRC, augments LGR5 expression independent of RAS/PIK3CA mutation status and promotes EGFR-LGR5 interactions. Furthermore, we describe the development of LGR5 ADCs incorporating a camptothecin-derived payload that is well-tolerated and significantly inhibits tumor growth. Importantly, cetuximab in combination with LGR5 ADCs results in enhanced tumor inhibition or regression versus single-agent treatment and extends survival in RAS^MUT^ patient-derived xenografts. These findings support growing evidence that ADC combination therapies may be more effective than monotherapies and suggests a broader clinical use for cetuximab in treating RAS^MUT^ CRC.

## Introduction

Leucine-rich repeat-containing G protein-coupled receptor 5 (LGR5) is a cell-surface seven-transmembrane receptor highly expressed in many cancers including liver, gastric, ovarian, and colorectal cancers (CRCs)^1–6^. LGR5 was initially identified as a marker of intestinal stem cells that gives rise to all other cell progenies in the crypt^7^. Our group and others showed LGR5 binds R-spondin 1-4 (RSPO1-4) ligands to potentiate Wnt/β-catenin signaling^8–10^. Further work identified LGR5 as a marker of CRC stem-like cells (CSCs) wherein ablation of adenomatous polyposis coli (APC), a negative regulator of Wnt signaling, in Lgr5^+^ stem cells resulted in the formation of intestinal adenomas^11^. Lgr5^+^ CSCs were shown to give rise to differentiated cell types (e.g., Lgr5^-^) while retaining long-term self-renewal capabilities^12,13^ and, importantly, elimination of Lgr5^+^ CSCs was shown to suppress primary tumor growth and reduce liver metastases^14^. Using mouse models and patient-derived data, others showed the majority of cancer cells disseminating from primary tumors to seed metastases were actually LGR5^-^; though reversion to an LGR5^+^ CSC state was required for metastatic colonization^15,16^. Furthermore, LGR5 has critical functions in mediating drug resistance as LGR5^+^ cells were shown to interconvert to a non-proliferative, drug-resistant LGR5^-^ state upon chemotherapy exposure^17,18^. Due to its overexpression in various cancers and critical roles in tumor initiation and progression, metastasis, and drug resistance, several attempts have been made to therapeutically target LGR5^1,19–21^.

Antibody-drug conjugates (ADCs) are one of the fastest growing classes of anti-cancer biologics and consist of highly specific monoclonal antibodies (mAbs) tethered to potent cytotoxic payloads via cleavable or non-cleavable chemical linkers^22–26^. Importantly, our group and others generated LGR5-targeting ADCs incorporating dipeptide enzyme-cleavable linkers attached to the microtubule inhibitor payload monomethyl auristatin E (MMAE)^1,19,20^ that showed tumor growth inhibition or regression in CRC xenograft models without notable toxicity^1,19^. However, a fraction of tumors relapsed following ADC withdrawal, likely due to target downregulation, suboptimal payload, and/or drug-resistant LGR5^-^ CRC cells that could evade treatment^1,17,18^. Another LGR5-targeting ADC with an acid-labile linker attached to the DNA-damaging anthracycline PNU159682 payload was shown to be too toxic to normal cells, including the intestinal crypts^19^. These findings suggest optimal safety and efficacy of ADCs is highly dependent on linker-payload selection. Further, an LGR5-targeting therapy alone may not be sufficient to eliminate colorectal tumors and may require a combinatorial strategy.

Epidermal growth factor receptor (EGFR) is overexpressed in 60-80% of CRCs, resulting in hyperactivation of KRAS and PI3K signaling pathways that promote proliferation and survival^27,28^, making it a favorable therapeutic target. Accordingly, there are several FDA-approved EGFR-targeting therapies including mAbs cetuximab (CTX) and panitumumab for RAS^WT^ metastatic CRCs (mCRCs), and tyrosine kinase inhibitors (TKIs), gefitinib, lapatinib, and osimertinib, among others, for different tumor indications^29–33^. However, resistance to these therapies is often observed, commonly through constitutively activating KRAS or PIK3CA mutations, thereby limiting the clinical benefit to patients without these mutations^27,34,35^. Interestingly, previous reports showed CTX treatment increased LGR5 mRNA levels in patient-derived tumor organoid (PDO) models of CRC^13,21^ and treatment with gefitinib increased the number of Lgr5^+^ cells in the mouse intestine^36^, suggesting EGFR inhibition (EGFRi) increases LGR5 expression. Consistently, LGR5 and other Wnt target genes were significantly upregulated in residual tumor cells from a panel of RAS^WT^ CRC patient-derived xenografts (PDXs) treated with CTX^37^. In line with these findings, CTX combined with LGR5 genetic ablation resulted in more significant tumor regression of CRC PDXs compared to treatment with CTX alone^13^. In this study, we examine EGFR-mediated regulation of LGR5 expression and evaluate the efficacy of EGFR-targeting therapies in combination with LGR5 ADCs for the improved treatment of CRC. We demonstrate CTX increases LGR5 protein levels irrespective of KRAS or PIK3CA mutations in preclinical models. We also identify a novel EGFR-LGR5 interaction that is augmented by CTX. Furthermore, we generated a camptothecin-based LGR5 ADC with a tripeptide linker (8E11-CPT2) utilizing site-specific, chemoenzymatic conjugation technology. Importantly, combination treatment of 8E11-CPT2 with CTX showed additive or synergistic activity in gastrointestinal (GI) cancer cells irrespective of mutational status and significantly reduced tumor burden and extended survival in RAS^MUT^ CRC PDX models compared to both monotherapies. Although CTX is only approved for RAS^WT^mCRC, these results suggest CTX administered in combination with LGR5 ADCs may feasibly expand its current clinical utility to mCRC patients with oncogenic mutations in the EGFR signaling pathway (e.g., KRAS or PIK3CA).

## Results

Inhibitors of the EGFR family increase LGR5 levels independent of KRAS or PIK3CA mutation status EGFR and LGR5 are highly expressed across GI cancer cell lines and PDX models with different RAS and PIK3CA mutation statuses as shown by western blot and RNA-seq data from the Cancer Cell Line Encyclopedia for models used throughout this study (Figure 1A-B). Specific gene mutations are listed in Table S1. As previous studies showed EGFRi increases LGR5 mRNA levels^13,21,36,37^, we first assessed the effect of inhibition of EGFR or the related and commonly targeted receptor HER2 on LGR5 protein expression in CRC cells with or without RAS^MUT^. LGR5-expressing LIM1215 (KRAS^WT^/PIK3CA^WT^), LoVo (KRAS^MUT^/PIK3CA^WT^), and SW620 (KRAS^MUT^/PIK3CA^WT^) CRC cells were treated with clinically-approved EGFR-targeting mAbs (CTX or nimotuzumab), HER2-targeting mAb (trastuzumab), or EGFR-(gefitinib) or EGFR/HER2-targeting (lapatinib) TKIs for 48 h. SW620 cells were used as a non-responsive control as they are negative for EGFR and express relatively low levels of HER2. EGFR-and, to a lesser extent, HER2-targeting therapies increased LGR5 protein levels in both LIM1215 and LoVo cells, but not in SW620 cells (Figure 1C), suggesting EGFRi-induced LGR5 upregulation occurs irrespective of KRAS mutation yet is dependent on EGFR expression.

**Figure 1.**
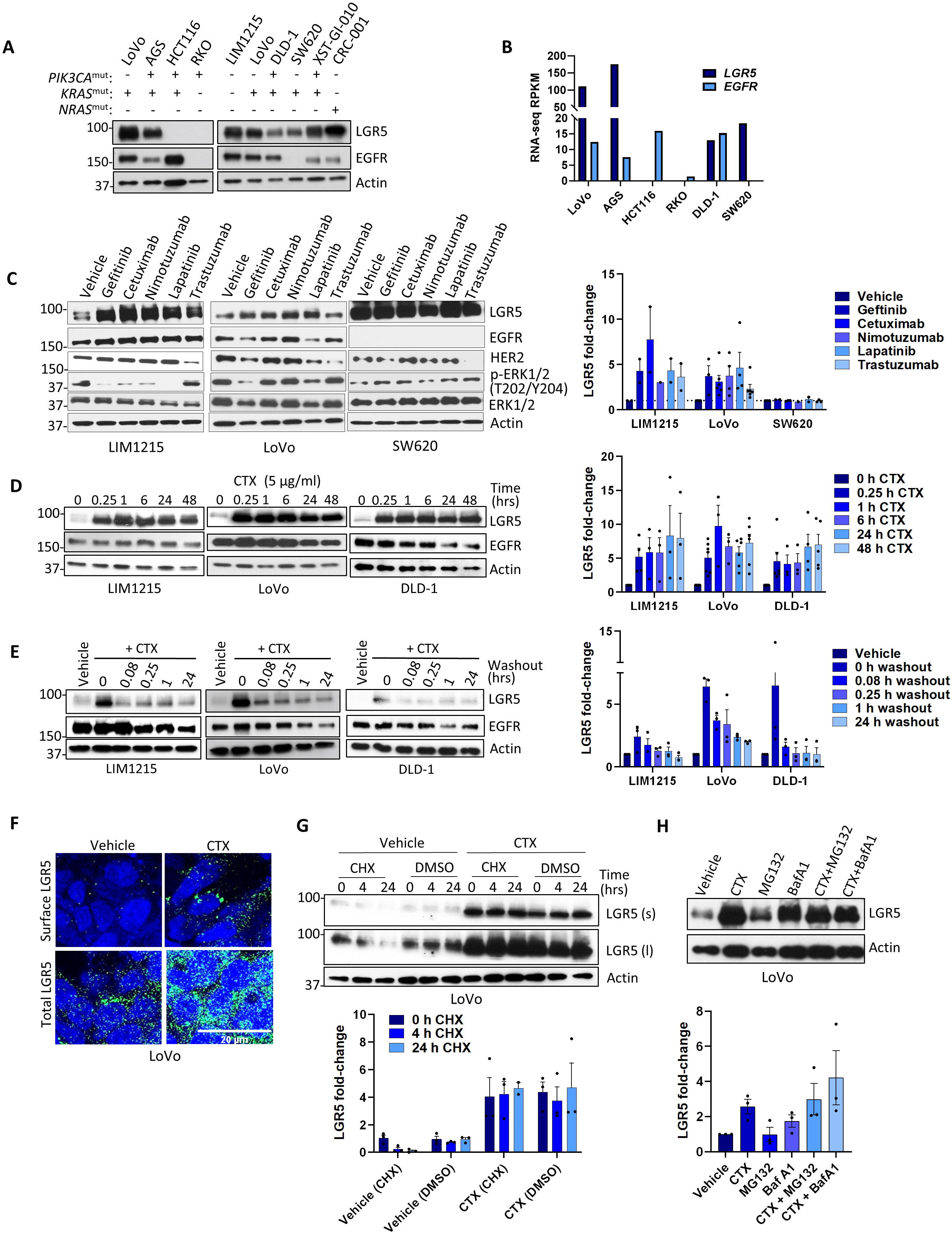
EGFR inhibitors reversibly increase LGR5 expression in CRC cells independent of mutation status. (A) Western blot of endogenous EGFR and LGR5 expression in a panel of CRC cell lines and patient-derived xenograft (PDX) models XST-GI-010 and CRC-001. (B) EGFR and LGR5 RNA-seq expression data from Cancer Cell Line Encyclopedia (CCLE). Values are read per kilobase of transcript per million (RPKM). Western blots and quantification of LGR5 protein fold-changes normalized to actin in (C) LIM1215, LoVo, and SW620 cells following 48 h treatment with a panel of EGFR-and HER2-targeting mAbs (3 µg/ml) and TKIs (1 µM), (D) 5 µg/ml CTX over indicated time-course, and (E) LIM1215, LoVo, and DLD-1 cells following 15 min CTX treatment and subsequent washout for indicated time-course. (F) Immunocytochemistry staining of total (permeabilized) and surface (non-permeabilized) LGR5 in LoVo cells treated for 15 min with vehicle or 5 µg/ml CTX. (G-H) Western blot and quantification of LGR5 protein fold-changes normalized to actin in (G) cycloheximide (CHX, 30 µg/ml) chase for 4 or 24 h performed in LoVo cells pre-treated for 15 min with vehicle or 5 µg/ml CTX (s, short exposure; l, long exposure) and (H) LoVo cells pre-treated for 15 min with 0.1 µM Bafilomycin A1 or 10 µM MG132 to inhibit lysosomal or proteasomal degradation, respectively, in the presence or absence of 5 µg/ml CTX.

We then focused primarily on CTX-mediated LGR5 upregulation for subsequent experiments as CTX is the only inhibitor from the panel approved for mCRC. Following 48 h treatment, CTX increased LGR5 in a dose-dependent fashion in LIM1215, LoVo, and DLD-1 (KRAS^MUT^/PIK3CA^MUT^) cells, demonstrating LGR5 upregulation occurs independent of both KRAS and PIK3CA mutations (Figure S1A). Furthermore, CTX-mediated LGR5 upregulation occurred within 15 min in all three cell lines (Figure 1D). Consistently, LGR5 upregulation was observed following 15 min treatment with trastuzumab or TKIs (Figure S1B). Trastuzumab was more effective at increasing LGR5 levels at the 15 min compared to TKIs, whereas the opposite was observed after 48 h treatment, likely due to different mechanisms of action (Figure 1C and S1B). Additionally, washout of CTX following 15 min treatment decreased LGR5 levels, suggesting CTX is required to sustain LGR5 upregulation (Figure 1E). Our findings were corroborated by immunocytochemistry (ICC), wherein 15 min CTX treatment promoted an increase in both total and surface levels of LGR5 (Figure 1F and S2A). In line with previous reports^13,21^, CTX significantly increased LGR5 mRNA levels in LIM1215 cells in a time-dependent fashion (Figure S2B). However, LGR5 mRNA remained relatively unchanged in LoVo cells (Figure S2C), suggesting LGR5 transcriptional upregulation may depend on KRAS mutational status. Given that LGR5 is a Wnt target gene^38^ and previous reports have shown both positive and negative regulatory roles of LGR5 in mediating Wnt/β-catenin signaling^8,18,39^, we tested the effect of CTX on activation of β-catenin. As shown in Figure S2D, CTX increased β-catenin activation in LIM1215 cells, but not LoVo cells, suggesting Wnt/β-catenin signaling is likely mediating transcriptional upregulation of LGR5 in LIM1215 cells. Notably, LoVo cells also harbor a mutation in APC. Taken together, these results suggest CTX and other EGFR-and HER2-targeting therapies increase LGR5 protein expression irrespective of KRAS or PIK3CA mutations, whereas activation of β-catenin and LGR5 transcription may be dependent on mutational status.

Typically, LGR5 is rapidly and constitutively internalized and degraded^40,41^. Therefore, one potential explanation for increased LGR5 protein levels in the absence of mRNA upregulation is LGR5 stabilization, which prevents or delays its internalization and subsequent turnover. To test this hypothesis, we examined the effect of CTX on LGR5 protein stability in LoVo cells by inhibiting de novo protein synthesis using a cycloheximide (CHX) chase assay. Remarkably, CTX stabilized LGR5 protein expression, whereas near-complete LGR5 degradation was observed for vehicle-treated cells after 24 h in the presence of CHX as shown for LoVo and LIM1215 cells (Figure 1G and S2E). To determine whether proteasome-or lysosome-mediated degradation is suppressed by CTX, we treated cells with proteasomal inhibitor MG132 or lysosomal inhibitor Bafilomycin A1 (BafA1) in the presence or absence of CTX. Interestingly, though basal LGR5 degradation was mostly suppressed by BafA1 rather than MG132, lysosomal inhibition had minimal effect after CTX treatment (Figure 1H and S2G). These results suggest upregulation of LGR5 by CTX likely occurs through post-translational stabilization.

### Loss of EGFR decreases LGR5 expression

We next examined how alternative perturbations of EGFR signaling affect LGR5 expression. Interestingly, siRNA-mediated knockdown (KD) of EGFR resulted in a concomitant reduction of LGR5 in LIM1215, LoVo, and DLD-1 cells (Figure 2A). EGF ligand-induced activation and subsequent downregulation of EGFR also reduced LGR5 protein levels in all three CRC cell lines (Figure 2B-C), consistent with a previous report that showed EGF reduced LGR5 in colorectal adenoma cells^42^. We next tested the effect of signaling inhibition downstream of EGFR after 48 h treatment with MEK1/2 inhibitor, trametinib (Figure 2D). Curiously, trametinib induced downregulation of EGFR and LGR5 in LoVo and DLD-1 cells in a dose-dependent manner. To determine if trametinib-mediated loss of EGFR and LGR5 is dependent on either receptor, we treated DLD-1 shLGR5 (LGR5 KD)^43^ and EGFR-negative SW620 cells with increasing concentrations of trametinib. As shown in Figure 2D, trametinib induced EGFR downregulation in LGR5 KD cells but did not promote loss of LGR5 in SW620 cells. This suggests trametinib-mediated EGFR downregulation is LGR5-independent, whereas loss of LGR5 is EGFR-dependent. Furthermore, loss of LGR5 occurred independent of ERK activation, as we observed no obvious decrease in LGR5 despite reduced ERK phosphorylation in SW620 cells. Notably, trametinib-mediated loss of LGR5 was not observed until 6 h with near complete loss at 24 h, suggesting downregulation may be mediated by transcriptional repression (Figure S3A). MEK1/2 inhibition by U0126 also resulted in loss of EGFR and LGR5 (Figure S3B). These results show loss of EGFR through divergent mechanisms results in a concomitant downregulation of LGR5.

**Figure 2.**
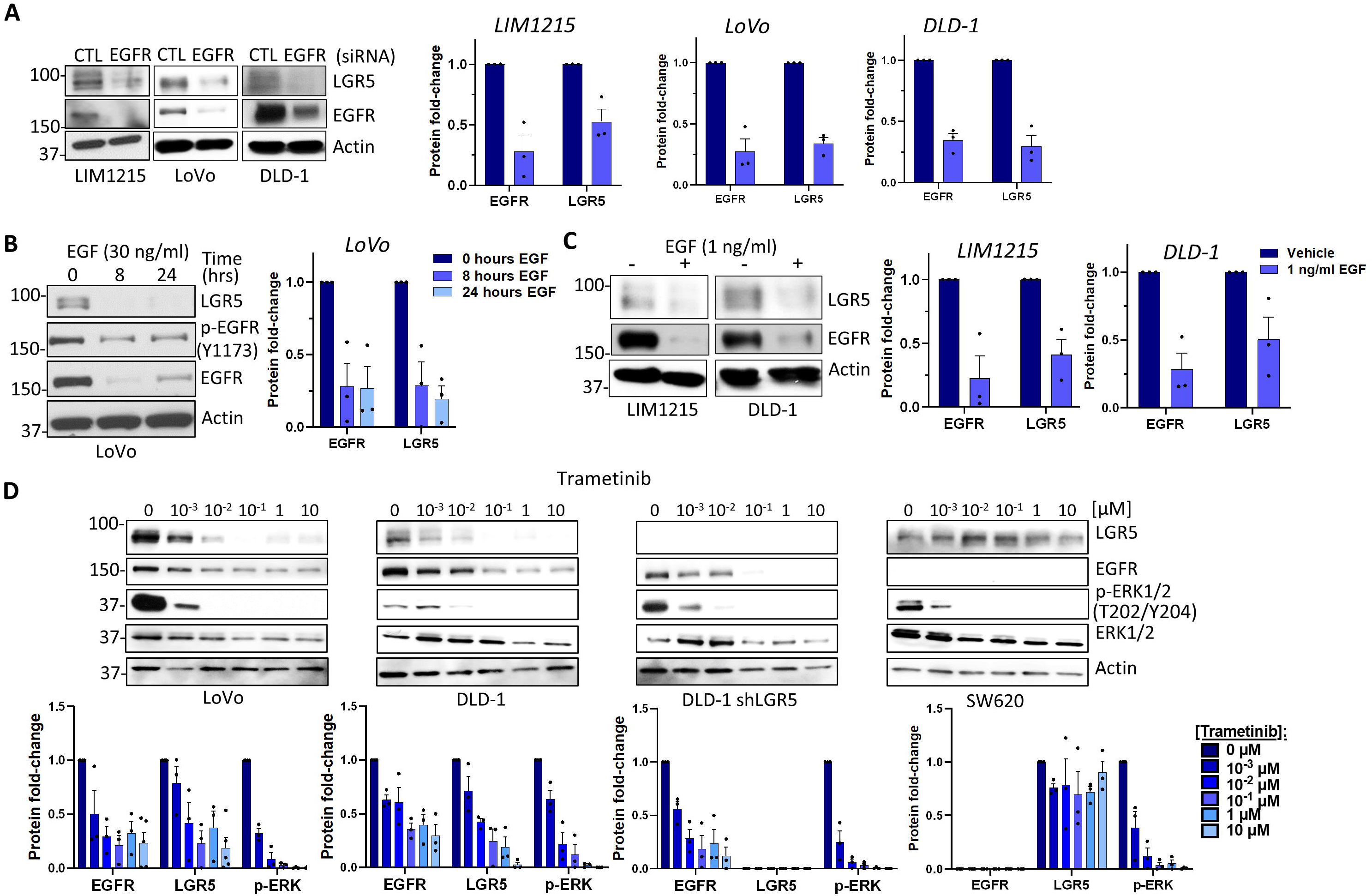
LGR5 levels are reduced simultaneously with loss of EGFR expression in CRC cells. Western blot and quantification following (A) 72 h treatment with 100 nM EGFR siRNA or non-targeting control siRNA in LIM1215, LoVo, and DLD-1 cells, (B) treatment with 30 ng/ml EGF for 8 or 24 h in LoVo cells, (C) 48 h treatment with 1 ng/ml EGF in LIM1215 and DLD-1 cells, and (D) 48 h treatment of LoVo, DLD-1, DLD-1 shLGR5, and SW620 cells with increasing doses of MEK1/2 inhibitor trametinib. EGFR and LGR5 protein fold-changes were normalized to actin and p-ERK/2 protein fold-change was normalized to total ERK1/2 for western blot quantification.

### EGFR interacts with LGR5 and is augmented by CTX

Having shown EGFR and LGR5 are co-degraded and CTX promotes stabilization of LGR5, we next examined whether EGFR and LGR5 interact. Co-immunoprecipitation (co-IP) experiments were performed using HEK293T cells overexpressing LGR5 (293T-LGR5)^8^ and LoVo cells to test for EGFR interaction with recombinant and endogenous LGR5, respectively. LGR5 co-IP’d with EGFR in both cell lines using an anti-EGFR antibody, but not the isotype control antibody (Figure 3A-B). To verify this interaction, we showed EGFR co-IP’d with LGR5 in LoVo cells using an anti-LGR5 mAb (Figure 3C). Interestingly, 15 min CTX treatment in LoVo cells dramatically increased the amount of LGR5 associated with EGFR, suggesting CTX promotes EGFR-LGR5 interaction (Figure 3D). To confirm these findings, proximity ligation assays (PLA) were performed in LIM1215 and LoVo cells. As expected, EGFR-LGR5 interactions significantly increased >2-fold after 15 min CTX treatment (Figure 3E-F). Together, these data demonstrate LGR5 interacts with EGFR and is augmented by CTX treatment.

**Figure 3.**
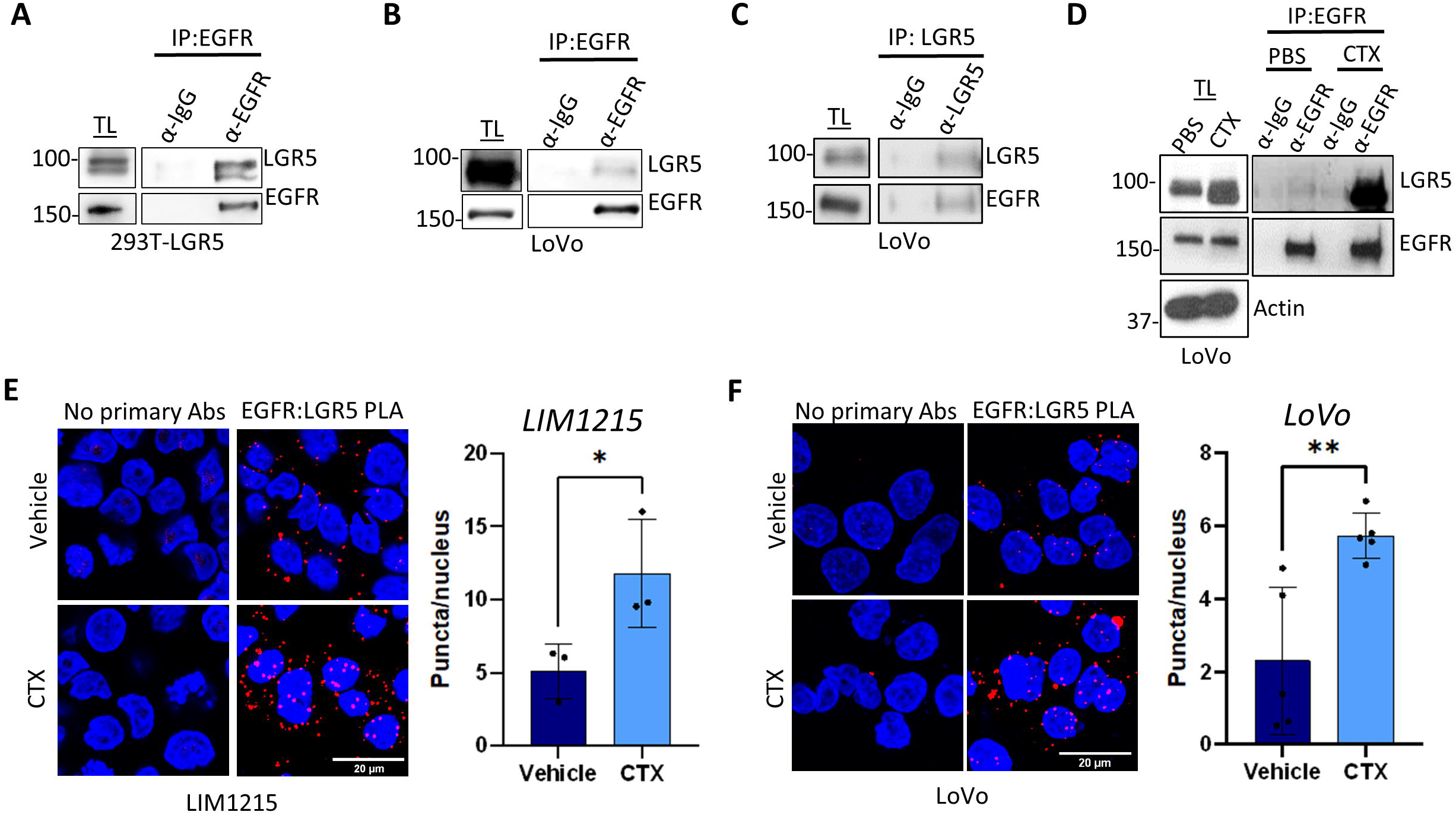
EGFR interaction with LGR5 is enhanced by CTX. Co-immunoprecipitation (co-IP) experiments with EGFR antibody or IgG isotype control shows (A) recombinant LGR5 in 293T-LGR5 cells and (B) endogenous LGR5 in LoVo cells specifically interacts with endogenous EGFR. (C) Co-IP experiment with LGR5 mAb in LoVo cells shows EGFR pulls down with endogenous LGR5. (D) Co-IP shows enhanced LGR5 interaction with EGFR in LoVo cells after 15 min treatment with 5 µg/ml CTX. TL, total lysate. Proximity ligation assays and quantification performed in (E) LIM1215 and (F) LoVo cells treated with vehicle or 5 µg/ml CTX. Statistical analysis was performed using two-tailed unpaired t-test; *p<0.05; **p<0.01. Data presented as mean +/-SD.

Generation and characterization of an LGR5-targeting ADC incorporating a camptothecin-derived payload Next, we generated a novel LGR5-targeting ADC incorporating a camptothecin (CPT)-derived payload. A valine-lysine-glycine (VKG) tripeptide linker attached to a glycine analog of 7-aminomethyl-10,11-methylenedioxy CPT (CPT2) was selected as the linker-payload due to its favorable safety profile, sensitivity of CRC to topoisomerase I inhibition, and increased potency compared to approved ADC linker-payloads deruxtecan and govitecan^44^. Notably, tripeptide linkers have shown to have increased efficacy and stability compared to more conventional dipeptide and acid-labile linkers^44–46^. First, we cloned, produced, and purified the LGR5-targeting 8E11 mAb with species cross-reactivity to both human and mouse LGR5 (K_d_=0.2 nM and 0.4 nM, respectively)^19^. 8E11 purity was confirmed by SDS-PAGE and Coomassie staining (Figure S4A). 8E11 binding to LGR5, internalization, and lysosomal trafficking, which is essential for ADC payload release, was verified by ICC and LAMP1 co-localization (Figure S4B). To develop the ADC, we performed site-specific conjugation utilizing microbial transglutaminase (mTG) and click chemistry to attach the payload to reactive glutamine (Q) residues in the mAb^47,48^. In addition to the Q295 residue located in the Fc region, we introduced an N297Q mutation to generate an 8E11 “QQ” mAb with two functional conjugation sites per heavy chain (Figure 4A). Compared to ADCs generated using stochastic lysine-or cysteine-based conjugation strategies, site-specific conjugation enables control over drug loading with improved ADC homogeneity and therapeutic index^49^. MTG catalyzed the addition of a branched, azido-functionalized spacer followed by strain-promoted azide-alkyne cycloaddition of a cleavable DBCO-functionalized PEG8-VKG-CPT2 (Figure 4A). Successful conjugation of two spacers and four linker-payload moieties per 8E11 heavy chain was confirmed by LC-MS/MS (drug-to-antibody ratio (DAR) = 7.94) and by SDS-PAGE and Coomassie staining (Figure 4B-C). A non-targeting, control ADC (R20-CPT2) incorporating the human B-cell CD20-targeting mAb (Rituximab®, R20) and identical linker-payload as 8E11-CPT2 was generated to assess off-target ADC cytotoxicity. R20 and R20-CPT2 purity were confirmed by Coomassie and successful linker-payload conjugation by LC-MS/MS (DAR = 8.00) (Figure S5A-C).

**Figure 4.**
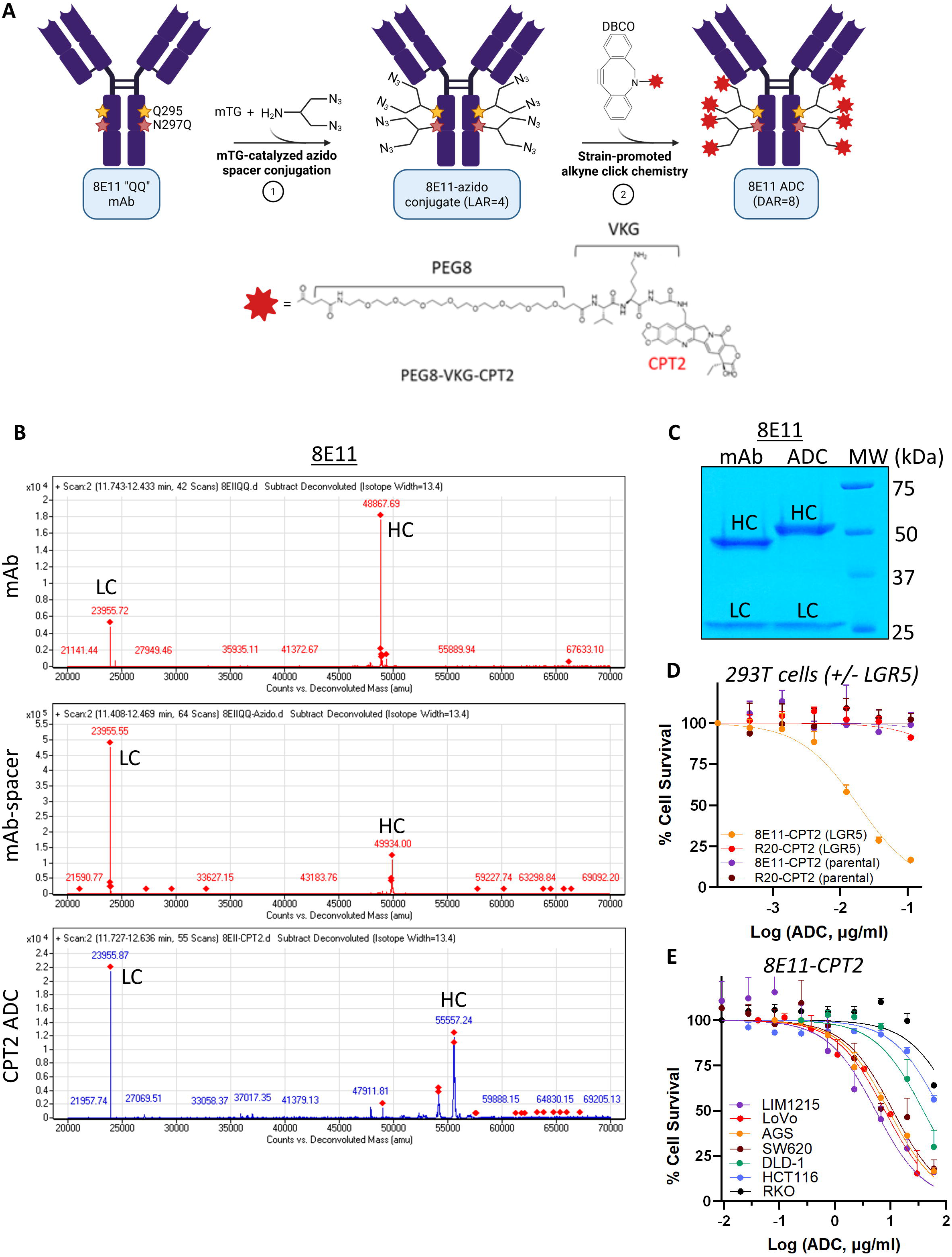
Generation and characterization of LGR5-targeting ADC 8E11-CPT2 (A) Schematic of microbial transglutaminase (mTG)-mediated conjugation of a branched linker to Q295 and N297Q of the Fc region of 8E11 mAb followed by strain promoted alkyne-azide cycloaddition of camptothecin 2 (CPT2) payload attached to a cleavable tripeptide valine-lysine-glycine (VKG) linker to generate 8E11-CPT2 ADC (DAR1=18). (B) LC-MS/MS analysis of 8E11 mAb, 8E11-azido spacer conjugate, and 8E11-CPT2 ADC under reducing conditions. LC, light chain; HC, heavy chain. (C) Coomassie staining of 8E11 mAb and 8E11-CPT2 ADC under reducing conditions shows increase in HC molecular weight, indicating linker-payload conjugation. (D) 8E11-CPT2 and R20-CPT2 cytotoxicity in LGR5-negative 293T parental and LGR5-overexpressing 293T-LGR5 cells. (E) Comparison of 8E11-CPT2 cytotoxicity in a panel of gastrointestinal cancer cell lines with different levels of LGR5 expression. Data presented as mean +/-SD.

### 8E11-CPT2 ADC exhibits high specificity and efficacy in vitro

Following the generation of 8E11-CPT2 and R20-CPT2 ADCs, we evaluated their specificity and cell-killing potency in vitro. To verify ADC target specificity, we evaluated 8E11-CPT2 cytotoxicity in 293T parental and 293T-LGR5 cells. 8E11-CPT2 exhibited high potency in 293T-LGR5 cells (IC_50_ = 0.032 +/-0.023 µg/ml) with minimal effects in LGR5-negative 293T cells (Figure 4D), demonstrating 8E11-CPT2 cell-killing was LGR5-dependent. R20-CPT2 showed negligible cytotoxicity in either cell line (Figure 4D). 8E11-CPT2 cytotoxicity was further assessed in a panel of GI cancer cell lines expressing different levels of LGR5 (Figure 1A-B and 4E). 8E11-CPT2 activity was correlated with LGR5 levels, exhibiting highest potencies in LGR5-high AGS gastric cancer cells and SW620 cells, followed by LoVo, LIM1215, and DLD-1 cells (Figure 4E). Minimal effects were observed in LGR5-negative HCT116 and RKO cells (Figure 4E). Average IC_50_ values are presented in Table S2. These data show 8E11-CPT2 mediates cell-killing efficacy in GI cancer cells expressing high levels of LGR5.

### Cetuximab enhances efficacy of LGR5-targeting ADCs in vitro

As CTX increases total LGR5 levels, we next evaluated if CTX treatment could enhance the efficacy of LGR5-targeting ADCs. We first confirmed that CTX pre-treatment would not negatively impact LGR5 mAb internalization and trafficking of mAb-LGR5 complexes to lysosomes in both LIM1215 and LoVo cells. Of note, rat LGR5 8F2 mAb was used for these experiments as 8E11 is humanized and CTX is a mouse/human chimera, both incorporating human IgG1 heavy chain and kappa constant regions. As shown in Figure 5A-B, CTX increased the amount of 8F2-LGR5 (∼2-fold) that trafficked to lysosomes. To evaluate combination treatment efficacy in vitro, LIM1215 and LoVo cells with high and low CTX sensitivity, respectively, and relatively resistant AGS and DLD-1 cells^50,51^ (Figure S6A) were treated with increasing concentrations of CTX and 8E11-CPT2 ADC. Drug synergy was assessed utilizing the Loewe additivity model which assumes the combination effect of two therapies to be equal to the sum of their normalized doses^52^ (Figure 5C-F). Typically, scores > 10 indicate a synergistic interaction,-10 to 10 an additive interaction, and values <-10 suggest an antagonistic interaction. In LIM1215 and AGS cells, combination treatment resulted in mainly additive cell-killing effects, approaching synergy at higher drug concentrations tested for each cell line. LIM1215 treatment with 0.05 µg/ml CTX or 20 µg/ml 8E11-CPT2 led to 47% and 45% cell death, respectively, whereas the combination resulted in 63% cell death (Figure 5G). AGS treatment with 20 µg/ml CTX or 6.67 µg/ml 8E11-CPT2 led to 28.8% and 28.4% cell death, respectively, whereas the combination resulted in 51.9% cell death (Figure 5H). However, more robust synergistic activity was observed in LoVo and DLD-1 cells. In LoVo cells, the combination of 0.25 µg/ml 8E11-CPT2 with 1 µg/ml CTX led to 48% cell death, whereas single-agent treatment with CTX or 8E11-CPT2 resulted in 31% or 12% cell death, respectively (Figure 5I). DLD-1 treatment with combination of 20 µg/ml 8E11-CPT2 with 10 µg/ml CTX led to 49% cell death, whereas single-agent treatment with CTX or 8E11-CPT2 resulted in 15% or 21% cell death, respectively (Figure 5J). Notably, CTX did not confer enhanced cell-killing with 8E11-CPT2 in EGFR-negative SW620 cells (Figure S6B). Loewe synergy scores for each tested combination are presented in Table S3. In line with these findings, CTX also enhanced the efficacy of our previously-reported LGR5-targeting ADCs incorporating MMAE and pyrrolobenzodiazepine dimer (PBD) payloads in LIM1215 and LoVo cells (Figure S6C-E)^1,53^. Taken together, these data suggest CTX enhances LGR5 ADC efficacy in vitro irrespective of ADC payload and KRAS mutation by increasing target expression, a critical determinant of ADC potency.

**Figure 5.**
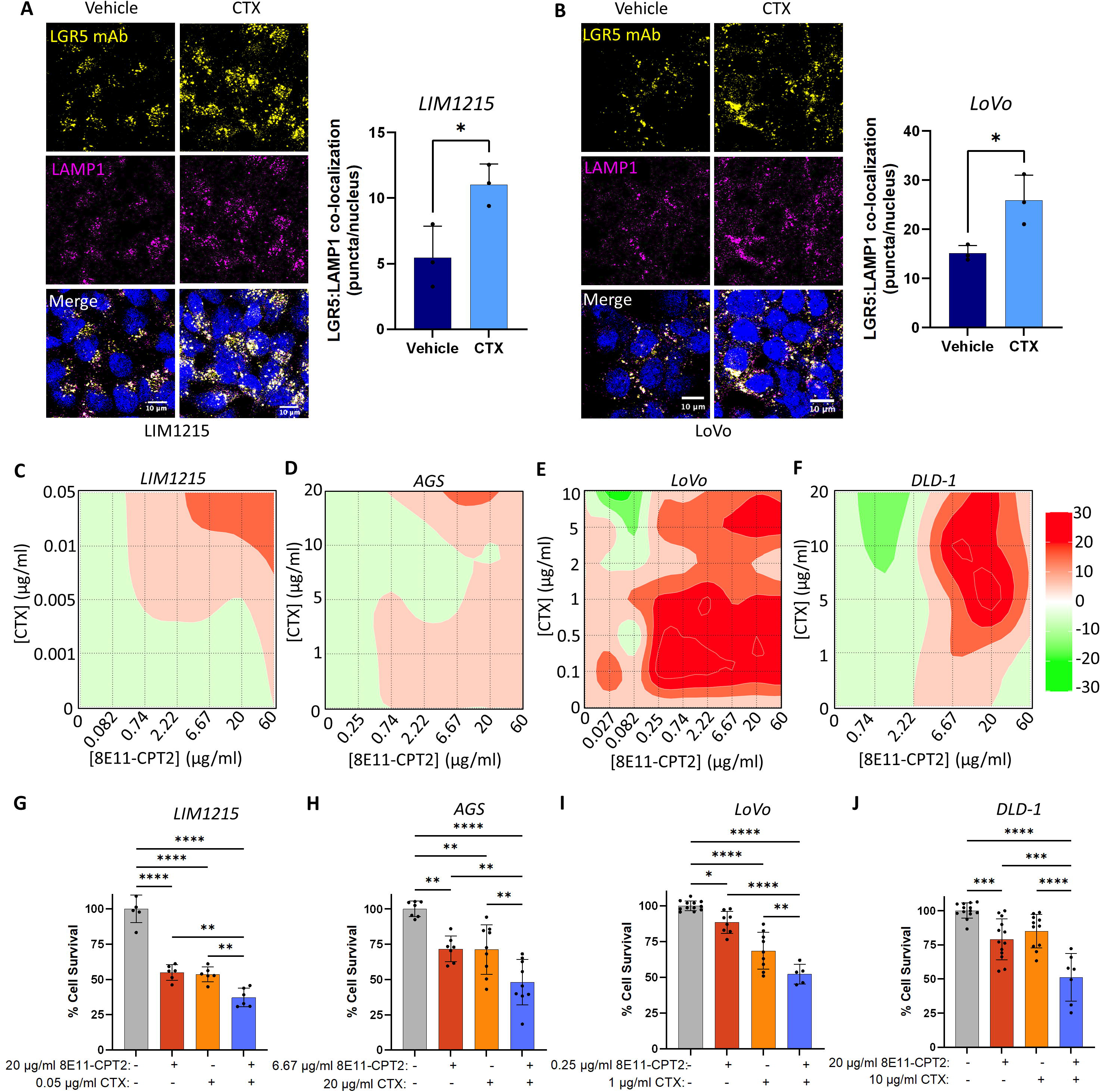
CTX enhances the efficacy of 8E11-CPT2 ADC in vitro. Confocal images and quantification of rat 8F2 LGR5 mAb internalization and co-localization with lysosome marker LAMP1 after 45 min incubation at 37° C in (A) LIM1215 and (B) LoVo cells pre-treated with vehicle or 5 µg/ml CTX for 15min. Statistical analysis was performed using two-tailed unpaired t-test; *p<0.05; **p<0.01. Data presented as mean +/-SD. Loewe synergy score heat maps and associated cell survival plots for (C) LIM1215, (D) AGS, (E) LoVo, and (F) DLD-1 cells co-treated with increasing concentrations of CTX and 8E11-CPT2 for 5 days. Example experiments showing % cell survival values for CTX and 8E11-CPT2 combinations with marked additive effects for (G) LIM1215 and (H) AGS cells and synergistic effects for (I) LoVo and (J) DLD-1 cells. Statistical significance was performed using ANOVA. *p<0.05; **p<0.01; ***p<0.001; ****p<0.0001.

Combination of cetuximab with 8E11-CPT2 results in enhanced antitumor efficacy and survival in patient-derived xenograft models of CRC To first determine if CTX can increase LGR5 levels in vivo, we tested increasing doses of CTX in a LoVo cell line-derived xenograft (CDX) model. Mice were administered 2 or 20 mg/kg CTX every 3 days for 2 doses and tumors were harvested on day 4. As shown in Figure 6A, CTX increased LGR5 expression in a dose-dependent manner in vivo. Further, DLD-1 xenograft tumors treated with two weekly doses of 10 mg/kg CTX harvested from a previously reported survival study^46^ showed increased LGR5 expression compared to vehicle (Figure 6B). Similarly, CTX increased LGR5 expression ex vivo in PDOs derived from the XST-GI-010 PDX model that harbors mutations in KRAS and PIK3CA (Figure 1A and 6C). To corroborate these findings, we analyzed the effect of CTX treatment on LGR5 mRNA expression using publicly-available microarray data from 21 CTX-responsive CRC PDX models^37^. Consistently, LGR5 levels were significantly upregulated following acute (24-72 h) and chronic (6 week) CTX treatment (Figure 6D). Notably, all PDX models were wild-type for RAS, BRAF, PIK3CA with the exception of one PDX that harbored a H1047R PIK3CA mutation. These data show CTX augments LGR5 expression levels in vivo.

**Figure 6.**
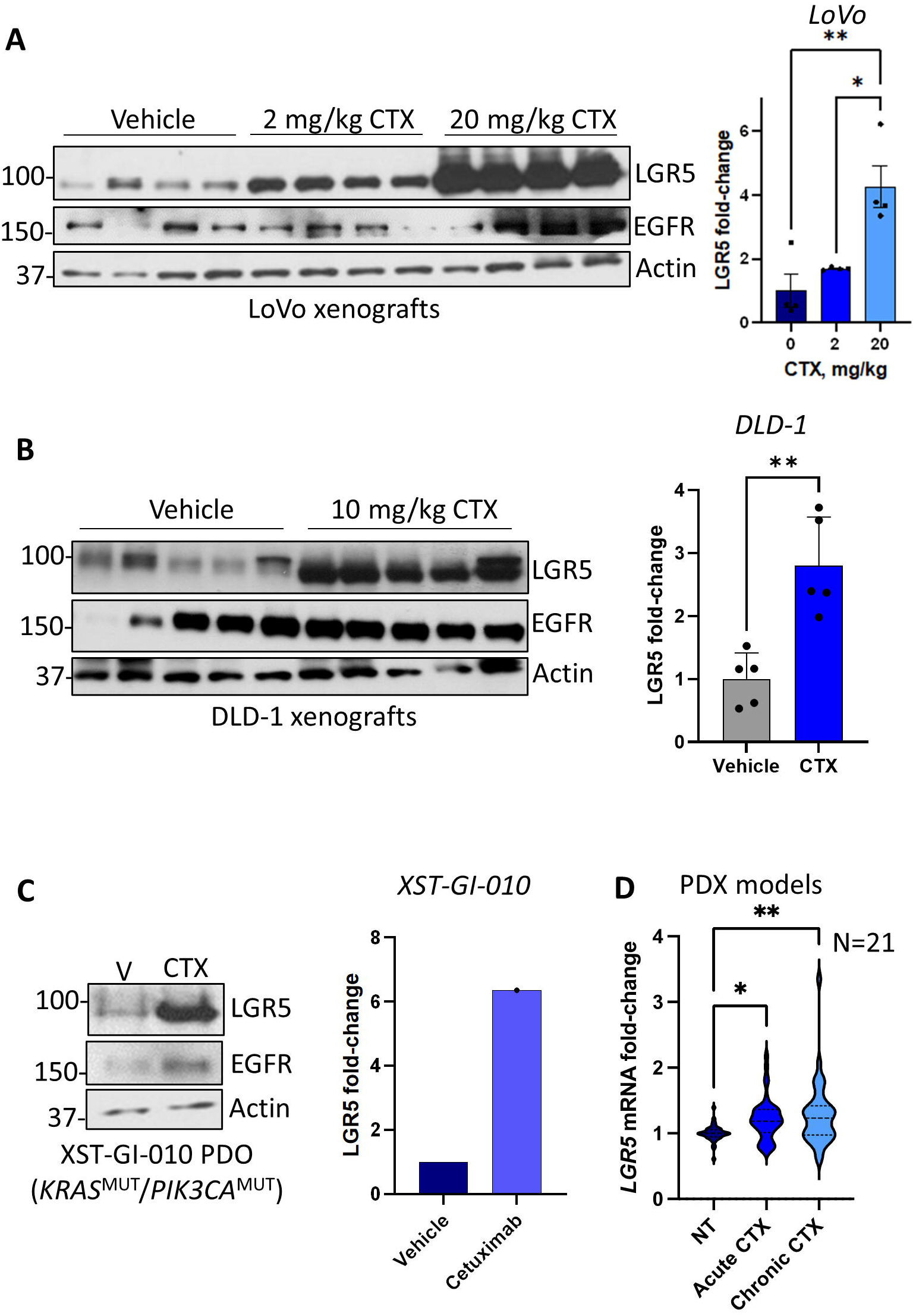
CTX treatment increases LGR5 expression in CRC models in vivo. Western blot and quantification of LGR5 expression in (A) LoVo cell line-derived xenograft models treated with 2 or 20 mg/kg CTX or PBS vehicle (N=4/group) once every three days for a total of two doses (tumors were harvested on day 4), (B) DLD-1 cell line-derived xenograft models from a previously reported survival study treated weekly for 2 doses of 10 mg/kg CTX or PBS vehicle (N=5/group), and (C) KRAS^MUT^ XST-GI-010 PDOs treated for 72 h with vehicle or 5 µg/ml CTX. LGR5 protein fold-change was normalized to actin. Data presented as mean +/-SD. (D) LGR5 mRNA microarray expression data following acute (24 to 72 h) or chronic (6-week) CTX treatment (20 mg/kg twice per week) in RAS^WT^ mCRC PDX models (GSE108277; N=21). Statistical analyses performed using ANOVA. *p<0.05; **p<0.01.

Prior to evaluating 8E11-CPT2 efficacy as a single agent or in combination with CTX in vivo, we conducted a safety study for 8E11-CPT2 in immunocompetent C57BL/6 mice. Mice were administered escalating doses of 8E11-CPT2 at 0, 5, 10, and 20 mg/kg and, after 2 weeks, blood was collected and hematology and clinical chemistry analyses were performed. Importantly, no significant changes in body weights, liver enzymes (alanine aminotransferase; ALT and aspartate aminotransferase; AST), kidney enzymes (creatinine; CRE), nor white blood cell (WBC) counts were observed up to 20 mg/kg (Figure 7A-E). Additionally, H&E staining of liver, kidneys, and intestines revealed no abnormal changes in tissue morphology (Figure S7-S8). These findings suggest 8E11-CPT2 ADC is well-tolerated with no observable toxicities.

**Figure 7.**
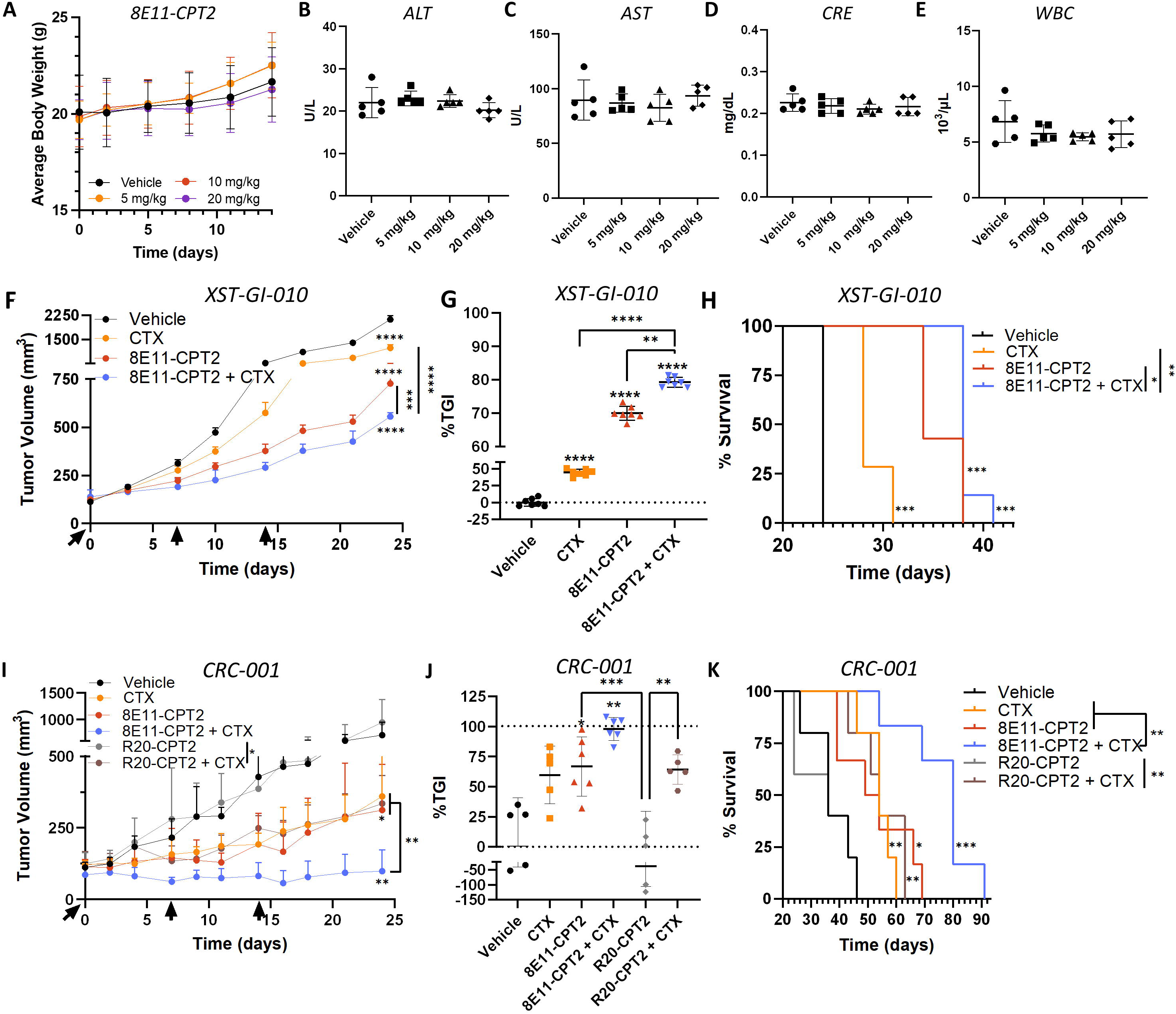
CTX enhances 8E11-CPT2 ADC antitumor efficacy and extends survival in patient-derived CRC models. Safety assessment shows (A) bodyweights, (B) ALT, (C) AST, (D) CRE, and (E) WBC plasma levels of C57BL/6J mice (N=5/group) following 2-week treatment with single-dose 8E11-CPT2 as indicated. (F) Anti-tumor efficacy of CTX or 8E11-CPT2 ADC compared to combination therapy in XST-GI-010 PDX models up to day 24 when first vehicle-treated animal reached maximal tumor burden. All treatments were performed at 5 mg/kg by intraperitoneal injection weekly for a total of 3 doses (N=7/group). Statistical significance performed using ANOVA. ***p<0.001; ****p<0.0001. Data presented as mean +/-SD. (G) Percent tumor growth inhibition (TGI) at day 24 in XST-GI-010 PDX models. Data presented as mean +/-SD. (H) Kaplan–Meier survival plot and log-rank test for the XST-GI-010 study. (I) Anti-tumor efficacy of CTX, 8E11-CPT2, and R20-CPT2 monotherapies and combination therapies in an NRAS^MUT^ CRC-001 PDX model up to day 24 when first vehicle-treated animal reached maximal tumor burden. All treatments were performed at 5 mg/kg by intraperitoneal injection weekly for a total of 3 doses (N=5 for all groups except 8E11-CPT2 and 8E11-CPT2+CTX, N=6). Statistical significance performed using ANOVA. *p<0.05; **p<0.01. Data presented as mean +/-SD. (J) TGI at day 24 in CRC-001 PDX models. Data presented as mean +/-SD. (K) Kaplan–Meier survival plot and log-rank test for the CRC-001 study.

We next evaluated the effects of CTX in combination with 8E11-CPT2 in LGR5-expressing, RAS^MUT^ XST-GI-010 and CRC-001 PDX models (Figure 1A). XST-GI-010 PDXs, derived from a cecal tumor, were treated with vehicle, CTX (5 mg/kg), 8E11-CPT2 (5 mg/kg), or CTX in combination with 8E11-CPT2 weekly for a total of 3 doses. CRC-001 PDXs, derived from a liver metastasis of a rectal tumor, were dosed similarly with additional treatment arms for R20-CPT2 (5 mg/kg) and R20-CPT2 in combination with CTX. Importantly, in both PDX models, CTX in combination with 8E11-CPT2 significantly inhibited tumor growth compared to all other treatment arms and conferred the greatest overall survival benefit (Figure 7F-K and S9). Notably, in the CRC-001 model, all tumors initially regressed with CTX/8E11-CPT2 combination with regrowth observed after treatment termination at days 14-21 (n=3, Figure S9B). However, other animals in the combination cohort showed continued regression until day 31 (n=2) or day 49 (n=1, Figure 7L-M and S9B). 8E11-CPT2 monotherapy showed slight regression in 3/6 animals before treatment termination (Figure S9B). No significant changes in body weight were observed between treatment groups (Figure S10A-B). Analysis of CRC-001 tumors after termination endpoints showed increased levels of LGR5 after CTX treatment, as expected, and in the majority of the CTX/8E11-CPT2 cohort examined, suggesting tumors would likely be responsive to additional LGR5-targeted therapy (Figure S10C). All tumors analyzed retained EGFR expression. These findings show 8E11-CPT2 ADC exhibits significant antitumor efficacy in LGR5-expressing, RAS^MUT^ CRC that can be further enhanced when administered in combination with CTX.

## Discussion

The majority of CRC-associated mortality arises from mCRC which has an approximate 14% 5-year survival rate^54^. Approximately 15-30% of patients present with mCRC at diagnosis whereas an additional 20-50% diagnosed with localized CRC eventually develop metastases^55^. In large part, the low survival rate of mCRC is due to intrinsic limitations of existing treatments. Approved therapeutic regimens consist mostly of combined fluorouracil and irinotecan chemotherapies with mAbs targeting vascular endothelial growth factor receptor (VEGFR; e.g. bevacizumab), EGFR (e.g. CTX or panitumumab), or programmed cell death protein 1 (PD-1; e.g. pembrolizumab and nivolumab)^56^. While chemotherapies may demonstrate initial efficacy, resistance and off-tumor toxicity (e.g. GI toxicity and neutropenia) commonly hinder sustained tumor regression^57,58^. Additionally, efficacy of EGFR-targeting mAbs is restricted to wildtype RAS and BRAF patients wherein only a modest survival benefit is observed in combination with chemotherapy^59^. Immune checkpoint blockade has proven efficacious in microsatellite instability-high and mismatch repair-deficient mCRCs, however these subtypes represent only a very small fraction (∼4-5%) of all mCRCs^60^. Furthermore, trastuzumab deruxtecan (T-DXd), a HER2-targeting ADC that has received tumor-agnostic approval for HER2-expressing solid tumors^61^, has shown antitumor activity in treatment-refractory, HER2-positive mCRC independent of RAS/PIK3CA mutations, although only 2-3% of mCRCs are HER2-amplified^62,63^. Thus, there is an urgent need to unveil new targets for improved mCRC therapies.

Despite the role of LGR5 in promoting tumorigenesis and disease progression, LGR5-targeting ADC strategies have thus far proven unsatisfactory^1,19^. Given previous works demonstrating EGFR inhibitors increase LGR5 expression^13,21,36,37^ and significantly reduce tumor size in combination with Lgr5 genetic ablation^13^, our work aimed to assess the antitumor efficacy of LGR5-targeted ADCs in combination with EGFR inhibitors, particularly CTX. Importantly, we showed CTX increased LGR5 protein levels independent of KRAS or PIK3CA mutation status and enhanced the efficacy of LGR5-targeted ADCs in several different LGR5-expressing preclinical models of CRC. These findings indicate that combination of CTX with other therapies, such as LGR5-targeted ADCs, may feasibly expand the CTX-responsive patient population beyond RAS^WT^ mCRCs.

Beyond therapeutic implications, this work also sheds important light on LGR5 signaling mechanisms and its potential crosstalk with the EGFR pathway. Presently, the functions of LGR5 in CRC remain elusive and contradictory^13–15,17,18,39,64–67^. Our work demonstrates that CTX induces LGR5 transcription in a cell line-dependent context, stabilizes LGR5 protein expression in the absence of de novo protein synthesis, and promotes EGFR-LGR5 interactions, raising several possible overlapping mechanisms for EGFR regulation of LGR5 expression. To our knowledge, this is the first work to report an interaction between EGFR and LGR5, though recent studies have reported interactions between EGFR and other members of the Wnt signaling complex including Fzd9b^68^, ZNRF3/RNF43 E3 ligases^69^, and LGR4^70^. Given EGFR-LGR4 interaction has been shown to promote breast cancer metastasis by reducing EGFR ubiquitination and degradation^70^, it would be interesting to examine the consequence of EGFR-LGR5 interaction on downstream signaling and ubiquitination of EGFR and LGR5 in future studies. Notably, while ZNRF3/RNF43 are known to interact with RSPO-LGR4 to inhibit Fzd clearance and potentiate Wnt/β-catenin signaling^71,72^, this interaction has not been observed with RSPO-LGR5, suggesting an alternative, unknown mechanism for LGR5. In contrast to EGFR inhibitors augmenting LGR5 levels, we show that when EGFR expression is decreased via EGFR KD or EGF-induced EGFR activation, LGR5 is downregulated. These findings suggest EGFR expression and its interaction with LGR5 is likely important for regulating LGR5 protein stability and degradation. Interestingly, we found treatment with high dose MEK inhibitors led to decreased EGFR and LGR5 levels after 24 h. We postulate that suppression of MEK/ERK signaling may result in transcriptional repression of EGFR and, in turn, loss of EGFR leads to LGR5 downregulation. However, MEK inhibition and/or EGF treatment may also decrease LGR5 at the transcriptional level, as the latter has been reported for adenoma cells^42^. Further, we observed that CTX upregulates LGR5 mRNA and induces β-catenin activation in KRAS^WT^ LIM1215 cells, but not KRAS^MUT^ LoVo cells, indicating that LGR5 transcriptional activation by CTX may be dependent on mutational status. As LGR5 is a known target gene of Wnt signaling^5^, it is possible that CTX induces β-catenin activation and subsequent Wnt signaling to induce LGR5 transcription in a KRAS-dependent fashion. However, further studies using additional CRC cell lines are necessary to validate these findings in order to fully understand EGFR-LGR5 signaling crosstalk and the relative contributions of transcriptional and post-translational mechanisms of LGR5 upregulation.

ADC efficacy in the clinic can often be limited by drug resistance and/or toxicity. Combination treatments with other systemic targeted therapies may therefore improve ADC efficacy and tolerability by lowering the threshold for an effective therapeutic dose^73^. In fact, EGFRi has been shown to increase efficacy of HER-targeting ADCs through a variety of mechanisms. For example, osimertinib, an EGFR TKI approved for T790M non-small-cell lung cancer, was shown to increase HER3 expression, improving internalization and efficacy of a HER3-DXd ADC^74^. Another study demonstrated CTX was able to overcome resistance to T-DXd mediated by EGFR overexpression and heterodimerization with HER2, which impaired HER2 internalization^75^. CTX treatment increased internalization of T-DXd bound to EGFR-HER2 heterodimers and improved ADC efficacy across several cancer types irrespective of KRAS mutation. This work suggests that CTX may also enhance ADC internalization, yet this is likely target-and tumor type-dependent. Given the enhanced efficacy of ADC combination therapies in preclinical studies, there are several ongoing clinical trials examining ADCs in combination with chemotherapy, endocrine therapy, radiotherapy, or other targeted treatments such as anti-angiogenic agents and immune checkpoint inhibitors^76^. For example, HER-targeted mAbs and TKIs have all been evaluated in combination with ADCs targeting HER2, TROP2, or MET for mCRC among other solid tumor indications^76^. CTX is currently being evaluated in combination with ADCs directed against different targets for solid tumor indications (e.g., ROR2; NCT05271604 and B7-H3; ARTEMIS-101). The aforementioned clinical trials coupled with our and others’ findings that CTX enhances the efficacy of LGR5-and HER2-targeting ADCs provide strong rationale for examining CTX in combination with ADCs in the clinic.

For reasons that remain poorly understood, KRAS mutation does not always confer resistance to CTX although it is likely alternative mediators are involved in determining CTX sensitivity^77–79^. Regardless, combination treatments of CTX with other targeted therapies represent one strategy to improve response to CTX and expand its clinical utility. For example, adagrasib, a G12C mutation-specific KRAS inhibitor, recently received FDA approval in combination with CTX based on results from the KRYSTAL-1 trial wherein combination with CTX promoted an increased median response duration compared to adagrasib alone^80,81^. Still, only ∼4% of mCRCs harbor KRAS G12C mutations^82^, emphasizing the need for mutation-agnostic combination treatments. In this work, we demonstrate LGR5 ADCs in combination with CTX results in more profound cancer cell-killing, tumor growth inhibition or regression, and survival compared to LGR5 ADC and CTX monotherapies irrespective of RAS mutational status. More broadly, EGFR and LGR5 dual-targeting approaches have shown clinical promise as demonstrated by petosemtamab, an EGFR:LGR5 bispecific antibody (bsAb) in phase II trials for mCRC (NCT03526835) and phase III trials for head and neck squamous cell carcinoma (NCT06496178; NCT06525220). Compared to CTX, petosemtamab showed increased inhibition of tumor growth and metastasis in KRAS^MUT^ CRC PDX models without affecting the expansion of normal colonic PDOs^21^. Interestingly, while both combination therapy and the bsAb approaches target EGFR and LGR5, the underlying therapeutic mechanisms of anti-tumor activity appear starkly different. Whereas petosemtamab specifically promotes internalization and degradation of EGFR in LGR5^+^ CRC cells, CTX and LGR5 ADC combination therapy relies on EGFR inhibition to increase LGR5 expression and thus enhance LGR5 ADC internalization and potency. It would be interesting, then, to directly compare the efficacy and safety profiles of combination treatments to petosemtamab in mCRC models. Overall, this study builds upon existing works to further our understanding of EGFR-LGR5 interactions and signaling as well as supports additional evaluation of EGFR and LGR5 dual-targeting approaches for treating RAS^WT^ and RAS^MUT^ mCRCs.

### Limitations of the study

While this study reveals a novel EGFR-LGR5 interaction, further work is necessary to fully understand the role of this interaction on downstream signaling and receptor stability. Unbiased approaches such as affinity purification mass spectrometry (AP-MS) and biotinylation identification (BioID) will help to resolve other proteins involved in this complex. Moreover, the presented work strongly supports further evaluation of EGFR and LGR5 dual-targeting approaches in the preclinical setting. 8E11-CPT2 was well-tolerated in mice up to the highest tested dose of 20 mg/kg without observed toxicities. Notably, another ADC with the same linker-payload was shown to be well-tolerated in rats at 60 mg/kg, given weekly four times^44^. As CTX does not recognize murine EGFR, we were unable to assess CTX toxicity in combination with 8E11-CPT2 although the 5 mg/kg CTX dose administered is far lower than recommended clinical doses (250–500 mg/m^2^). Further, LGR5 is expressed at much lower levels in normal tissues and LGR5-positive adult stem cells in the gut have shown to be dispensable, as they can be replaced by a reserved stem cell pool^83–85^. This suggests, as with most ADCs, any observed toxicity from 8E11-CPT2 would likely be payload-mediated rather than target-based. Though ADCs are routinely first tested in rodents, we should point out that topoisomerase I inhibitors are generally better tolerated in rodents, particularly mice, compared to human and non-human primates (NHPs). However, first-in-human dosing of novel ADCs can potentially be scaled to other clinical ADCs with the same or similar payloads^86^. As ADCs incorporating topoisomerase I inhibitor payloads such as T-DXd or datopotamab deruxtecan are tolerated at clinical doses of 5.4 or 6.4 mg/kg and 8 mg/kg^86^, respectively, this is why we selected 5 mg/kg for initial efficacy studies. However, alternative dosing concentrations and frequencies of CTX and 8E11-CPT2 ought to be evaluated in mice to potentially improve tumor regression and survival while minimizing any potential toxicities. With the goal of developing a more effective treatment for mCRC, this study focused on establishing proof-of-principle that our CTX/LGR5 ADC combination was more effective compared to CTX monotherapy in subcutaneous models, including CRC-001 derived from a liver metastasis of rectal cancer. However, future studies should also evaluate 8E11-CPT2 and CTX combination treatment in orthotopic mCRC models of different mutational backgrounds and molecular subtypes as well as test the combination treatment at in NHP safety studies. Ultimately, this study provides rationale for further evaluation of LGR5 ADCs in combination with CTX to improve the treatment of mCRC.

## RESOURCE AVAILABILITY

### Lead contact

Requests for resources and reagents should be directed to the lead contact, Kendra S. Carmon.

### Materials availability

All materials generated in this study are available through the lead contact.

### Data and code availability

- All raw data generated in this study are available upon request from the lead contact.
- Publicly available data can be found at Gene Expression Omnibus GSE108277 and Cancer Cell Line Encyclopedia (CCLE; www.cbioportal.org)
- This paper does not report original code.
- Any additional information required to reanalyze the data reported in this paper is available from the lead contact upon request.

## ACKNOWLEDGEMENTS

This work was supported by funding from NIH/NCI (R01 CA226894, R21 CA270716, and R21 CA282378) to K.S.C., and a predoctoral fellowship of the Gulf Coast Consortia, on the Training Interdisciplinary Pharmacology Scientists Program (T32 GM139801) to P.H. We would like to thank Dr. Vihang Narkar for use of equipment, Dr. Zhengmei Mao and Stephen Farmer for confocal microscopy technical assistance, Dr. Julie Rowe, Betty Arceneaux, and Dr. Karan Saluja with assistance in patient sample collection and processing, and Martha Thompson for assistance with regulatory approvals for research involving human subjects. Additionally, we would like to thank Dr. Sheng Pan and Li Li from the Clinical and Translational Proteomics Service Center at UTHealth Houston.

## AUTHOR CONTRIBUTIONS

Conceptualization and Design: K.S.C. and P.C.H.; Methodology: K.S.C., P.C.H., Y.T.; Investigation: K.S.C., P.C.H., Z.L., C.G., S.S., Y.M.S., A.M.A., Y.T.; Data curation: K.S.C., P.C.H., Z.L., C.G., S.S., Y.M.S, A.M.A., Y.T.; Formal Analysis: K.S.C., P.C.H., Y.T.; Writing original draft: P.C.H.; Review and Editing: K.S.C, P.C.H., C.G., Y.T.; Resources: K.S.C.; Funding acquisition: K.S.C..; Supervision: K.S.C.

## METHOD DETAILS

### Plasmids and cloning

The LGR5 mAb 8E11^19^ was generated from patented sequences and cetuximab (CTX), trastuzumab, and rituximab (R20) mAbs were generated utilizing publicly-available sequences. The variable heavy and light chain (VH and VL) regions were synthesized (Epoch Life Science, Inc.) and subcloned into pCEP4 vector containing either human IgG1 or kappa constant regions using In-Fusion Snap Assembly (Takara Cat# 638947) as we previously reported^87^. For 8E11 and R20, an N297Q mutation was introduced in the IgG heavy chain Fc region for site-specific conjugation.

### Cell culture

DLD-1 (Cat# CCL-221, RRID:CVCL_0248), LoVo (Cat# CCL-229, RRID:CVCL_0399), RKO (Cat# CRL-2577, RRID:CVCL_0504), HCT116 (Cat# CCL-247, RRID:CVCL_0291), COLO320 (CCL-220, RRID:CVCL_0219), SW620 (Cat# CCL-227, RRID:CVCL_0547), AGS (Cat# CRL-1739, (RRID:CVCL_0139), and HEK293T/293T

(Cat# CRL-3216, RRID:CVCL_0063) cells were purchased from ATCC. LIM1215 (Cat# 10092301, RRID:CVCL_2574) were purchased from Millipore Sigma. Cell lines were authenticated utilizing short tandem repeat profiling and tested for mycoplasma. 293Ts were cultured in DMEM and cancer cell lines in RPMI medium supplemented with 10% fetal bovine serum (Gibco Cat# 26140079) and penicillin/streptomycin (Gibco Cat# 15140122). LIM1215 cells were additionally supplemented with 1 µg/ml hydrocortisone (Sigma-Aldrich Cat# H0888), 10 µM 1-Thioglycerol (Sigma-Aldrich Cat# 88640), 0.6 µg/ml insulin (Gibco Cat# 12585014) and 25 mM HEPES (Gibco Cat# 15630080). Cell lines were cultured at 37°C with 95% humidity and 5% CO_2_. LoVo and DLD-1 LGR5 shRNA KD cell lines and stable 293T-LGR5 cell lines were generated as previously described and cultured in 1 µg/ml puromycin^1,8^ (Gibco Cat# 1113803).

### Patient-derived tumor organoid culture

To generate XST-GI-010 organoids, tumor tissue was cut into small pieces, washed with ice-cold PBS, and subsequently digested with Liberase TH grade (Roche Life Science Cat# 5401135001) for 11h at 371°C with vigorous pipetting every 15 min. The remaining fragments were treated with TrypLE Express (Gibco Cat# 12563011) at 371°C for 201min to disperse into single cells. The supernatant was collected and centrifuged at 200×g for 31min at 41°C. The cell pellet was suspended with growth factor reduced Matrigel (Corning Inc. Cat# 356231) and dispensed into 48-well culture plates (25 μl Matrigel/well). After Matrigel polymerization, complete medium (advanced DMEM/F12 supplemented with 50 ng/ml EGF (Gibco Cat# PMG8043), 100 ng/ml Noggin (PeproTech Cat# P97466), 500 nM A 83-01 (StemCell Tech. Cat# 72024), 10 nM Gastrin (Sigma-Aldrich Cat# G9145), 1× B-27 (Gibco Cat# 17504044), 11mM N-acetylcysteine (Thermo Scientific Cat# A1540914), 101mM HEPES, 21mM Glutamax (Gibco Cat# 35050061) and penicillin/streptomycin) was added and replenished every 2–3 days.

### Western blot

Protein lysate was prepared using RIPA lysis buffer (ThermoFisher Cat# 89901) supplemented with Halt protease and phosphatase inhibitors (ThermoFisher Cat# 78443). PDX tissues were homogenized and freeze-thawed. Lysates were sonicated then centrifuged 10 min, 4°C at 13,000xg. Protein concentrations were quantified by BCA protein assay (Thermo Scientific Cat# 23227) and lysates were diluted in reducing SDS Laemmli buffer (Thermo Scientific Cat# J60015.AD) and incubated at 37°C for 1 h prior to SDS-PAGE. Anti-mouse IgG (Cat# 7076, RRID:AB_330924) and anti-rabbit IgG (Cat# 7074, RRID:AB_2099233) HRP-labeled secondary antibodies from Cell Signaling Technology were utilized with the standard ECL protocol (Cytiva RPN3004). Primary antibodies used in this study include: anti-EGFR (Cat# 54359, RRID:AB_2799458, 1:1000); anti-phospho-EGFR Y1068 (Cat# 3777, RRID:AB_2096270, 1:1000)); anti-HER2 (Cat# 4290, RRID:AB_10557104, 1:1000); anti-β-actin (Cat# 3700, RRID:AB_2242334, 1:5000); anti-p44/42 MAPK/ERK1/2 (Cat# 9102, RRID:AB_330744, 1:1000); anti-phospho-p44/42 MAPK T202/Y204 p-ERK 1/2 (Cat# 9101, RRID:AB_331646, 1:1000); anti-β-catenin (Cat# 8480, RRID:AB_11127855, 1:1000); anti-non-phospho (active) β-catenin S33/37/T41 (Cat# 8814, RRID:AB_11127203, 1:1000) from Cell Signaling Technology and anti-LGR5 (Cat# ab75732, RRID:AB_1310281, 1:1000) from Abcam. Chemical inhibitors, ligands, and siRNA used include: gefitinib (Cat# HY-50895); lapatinib (Cat# GW572016); nimotuzumab (Cat# HY-P9968); and U0126 (Cat# HY-12031A) from MedChemExpress; trametinib (Selleck Cat# GSK1120212); CHX (Selleck Cat#S7418); EGF (Gibco Cat# PMG8043); EGFR siRNA (Cell Signaling Technology Cat# 6480); MISSION® negative control siRNA (Sigma Cat# SIC001). Western Blot quantification was performed with Fiji ImageJ by normalizing protein band intensity to the explicated reference protein (e.g., actin, ERK1/2, β-catenin). Relative protein-fold changes were calculated by dividing this value for each treatment condition by that of vehicle control.

### RT-qPCR

Total RNA was isolated using TRIzol Reagent (Thermo Fisher Cat# 15596026) followed by purification using the RNeasy Mini Kit (Qiagen Cat# 74104). cDNA was synthesized using SuperScript IV VILO master mix with ezDNase enzyme (ThermoFisher Cat# 11766050). RT-qPCR analysis was performed using amfiSure qGreen Master Mix (GenDEPOT Cat# Q5600-005) on CFX96 Touch Real-Time PCR Detection System (Bio-Rad). Each sample was run in triplicate. LGR5 mRNA was normalized to GAPDH and relative expression was calculated using the 2^(−ΔΔ^C ^)^ method. LGR5 primer sequences were (forward)-ATCTCATCTCTTCCTCAAA and (reverse)-CTTCTAATAGGTTGTAAGACA and, for GAPDH, (forward)-TCAAGGCTGAGAACGGGAAG and (reverse)-CGCCCCACTTGATTTTGGAG.

### Immunocytochemistry

Cells were seeded in a poly-D-lysine-coated 8-well chamber slide (Corning Cat# 354632) and treated the following day. For lysosome co-localization experiments, cells were treated with 5 µg/ml LGR5 mAb (8E11 or rat anti-LGR5 (8F2); BD Biosciences Cat# 748842, RRID:AB_2873245) at 37°C for 1 h with or without CTX, washed, fixed in 4% paraformaldehyde (ThermoFisher Scientific 28906) and permeabilized in 0.1% saponin (Sigma-Aldrich Cat# 84510). Cells were incubated with anti-LAMP1 (Cell Signaling Technology Cat# 9091, RRID:AB_2687579, 1:400) at room temperature for 1 h, followed by anti-human-Alexa-488 (ThermoFisher Cat# A-11013, RRID:AB_2534080) or anti-rat-Alexa-488 (ThermoFisher Cat# A-21210, RRID:AB_2535796) and goat anti-rabbit-Alexa-555 (ThermoFisher Cat# A-21428, RRID:AB_141784, 1:200) at room temperature for 1 h. To detect total versus surface levels of LGR5 following CTX treatment, cells were fixed and permeabilized before or after 1 h incubation with 5 µg/ml mouse anti-LGR5 (rm8F2) mAb^1^. Nuclei were counterstained with TO-PRO3 (1:1000; ThermoFisher Cat# T3605) for 15 min. Images were acquired using confocal microscopy (Leica TCS SP5, RRID:SCR_020233) and LAS AF Lite software. Images were quantified using ImageJ by normalizing total puncta to total nuclei in non-overlapping fields of >70 cells.

### Co-immunoprecipitation (co-IP)

Protein lysate was harvested using IP Lysis buffer (Pierce Cat# 87787) supplemented with Halt protease and phosphatase inhibitors. Lysates were sonicated then centrifuged 10 min, 4 °C at 13,000xg. For EGFR co-IP, EGFR mAb-conjugated beads (Cell Signaling Technology Cat# 5735, RRID:AB_10691854) or negative control isotype IgG-conjugated beads (Cell Signaling Technology Cat# 3420, RRID:AB_1549744) were added to lysate at 1:20 dilution and rotated overnight at 4°C. For LGR5 co-IP, 25 µl of Protein A magnetic beads (Pierce Cat# 88845) and 2 µg of rat anti-LGR5 (8F2) mAb (BD Biosciences Cat# 748842, RRID:AB_2873245) or rat IgG control (ThermoFisher Cat# 14-4031-82) were added to 150 µl protein lysate. The following day, samples were washed thrice with IP Lysis buffer, resuspended in 20 µL SDS Laemmli buffer, and heated at 50°C for 30 min prior to running SDS-PAGE. Primary and secondary antibody and ECL protocols were followed as previously described.

### Proximity ligation assay (PLA)

PLA was performed utilizing the DuoLink® kit (Sigma-Aldrich Cat# DUO92101). Briefly, cells were seeded overnight in 8-well chamber slides at approximately 75% confluency and treated the following day with CTX (5 µg/ml, 15 min). Cells were fixed and permeabilized as previously described, blocked for 1 h at 37°C, then incubated 1:200 with primary rabbit anti-EGFR (Cell Signaling Technology Cat# 54359, RRID:AB_2799458) and mouse anti-LGR5 (8F2)^1^ at 37°C for one hour. Cells were then treated with PLUS/MINUS PLA probes for one hour, ligase for 30 min, then polymerase for 100 min, all at 37°C. Finally, the slides were mounted with DAPI-containing mounting media and imaged the following day on a Nikon AX/AX R confocal microscope. Quantification was performed in Fiji ImageJ by normalizing total puncta to total nuclei in non-overlapping image fields of ≥70 nuclei.

### Antibody production

MAb production was performed by transient expression of corresponding light and heavy chain constructs in Expi293F cells (ThermoFisher Cat# A14527, RRID:CVCL_D615) using Polyethylenimine HCL Max reagent (Polysciences Cat# 247651). Media was collected 7 days post-transfection and mAbs were purified using protein A (Genscript Cat# L00210) affinity chromatography and eluted under acidic conditions (pH 3 glycine) into pH 8.5 TRIS. Buffer exchange to PBS was performed by centrifugation at 4000 rpm. MAbs were analyzed for purity and homogeneity by SDS-PAGE and LC-MS/MS with quatification by NanoDrop (ThermoFisher).

### Microbial transglutaminase (mTG) site-specific conjugation

For site-specific conjugation to Q295 and Q297, 8E11 mAb (6.50 mL in PBS; 5.016 mg/ml; 32.60 mg) was incubated with diazido branched linker, N-(Amino-PEG2)-N-bis(PEG3-azide) (BroadPharm Cat# BP-24369) (188 µL in H_2_O; 100 mg/mL; 160 equiv.) and Activa TI^®^ Transglutaminase (401 mg; final concentration 15%, Ajinomoto, Modernist Pantry) at RT for 16 h. Conjugated mAb was purified by protein A column chromatography to afford a mAb-linker conjugate [24.84 mg; 76% yield determined by Nanodrop]. R20 was conjugated similarly (74% yield).

### Strain-promoted azide-alkyne cycloaddition

DBCO-PEG8-VKG-CPT2 was synthesized (MedChemExpress) and added (139 µL of 20 mg/ml in DMSO, 1.5 equivalent per azide) to a solution of 8E11-linker conjugate (1.86 mL in PBS; 13.356 mg/ml; 24.84 mg), and the mixture was incubated at RT for 16 h. Crude products were purified by FPLC (Cytiva AKTA pure) to afford the 8E11-CPT2 ADC (5.711 mg/ml in PBS; 13.99 mg; 56.3% yield). Drug-antibody ratio (DAR) values were determined by LC-MS/MS analysis (DAR_8E11-CPT2_=7.94). R20-CPT2 ADC was generated and purified similarly (DAR_R20-CPT2_=8.00).

### Mass Spectrometry

MAb and ADC samples were analyzed at the Clinical and Translational Proteomics Service Center at UTHealth Houston on an Agilent 6538 UHD Accurate-Mass Quadrupole Time-of–Flight (Q-TOF) LC/MS system coupled with an Agilent 1200 series HPLC. LC separation was carried out on an Agilent PLRP-S reversed phase column (2.1 × 50 mm, 5µm, 1000Å,). The solvents were 2 % acetonitrile, 98% water, and 0.1% formic acid (A); 80% acetonitrile, 20% water, and 0.1% formic acid (B). A gradient of 25%-90% B was applied over 20 min at a flow rate of 0.2 mL/min. The samples were injected after being diluted with 25% acetonitrile, 75% water, and 0.1% formic acid. The Q-TOF was operated in ESI positive mode with capillary voltage 3500 V, drying gas flow rate of 7L/min and the source temperature of 325°C. Spectra were acquired in MS1 scan from 4.5-20 min each run over the mass range of 650-2800 m/z. Raw data were processed using Agilent MassHunter BioConfirm software (Version B.04.00) utilizing the Maximum Entropy deconvolution algorithm for the accurate molecular mass calculation; deconvoluted mass range was set at 20-100 kDa.

### In vitro cytotoxicity assays

CRC cells were plated at approximately 1,500 cells/well in 96 half-well plates (Corning Cat# 3885). Serial dilutions of 8E11-CPT2 or R20-CPT2 were added with or without CTX and incubated at 37°C for 4 or 5 days (all cell lines except LIM1215 were treated for 4 days). Cell viability was measured using the CellTiter-Glo 2.0 Assay (Promega Cat# G9242) according to the manufacturer’s protocol. Luminescence was measured using Tecan Infinite M1000 plate reader (RRID:SCR_025732). Synergy was calculated using the Loewe Additivity Model in SynergyFinder Plus^88^.

### Tolerability study

Female 6-8-week-old C57BL/6J mice (n= 5/group, The Jackson Laboratory, RRID:IMSR_JAX:000664) received a single dose 8E11-CPT2 (5, 10, or 20 mg/kg) by intraperitoneal injection. Body weight was monitored every 2-3 days for 2 weeks. Humane endpoints were defined as >20% weight loss or severe signs of distress. However, no mice met these criteria throughout the study. Fourteen days post-injection, mice were terminally anesthetized and whole blood was drawn by heart puncture for blood chemistry analysis in the Department Veterinary Medicine & Surgery at MD Anderson Cancer Center. Liver and kidney tissues were collected and then paraffin embedded, sectioned, and H&E stained in the histopathology core at UTHealth Houston Institute of Molecular Medicine.

### In vivo efficacy studies

Animal studies were carried out with IACUC approvals (AWC-20-0144, AWC-21-0142, and AWC-23-0106). For LoVo xenograft studies, nu/nu (Charles River Laboratories, RRID:IMSR_CRL:088) mice were implanted subcutaneously with approximately 2.5 x 10^6^ cells. When tumors reached 8-9mm in diameter, mice were treated with vehicle, 2 mg/kg CTX, or 20 mg/kg CTX at day 0 and day 3. Mice were euthanized and tumors collected at day 4 for analysis. For ADC efficacy studies in PDX models, NSG (The Jackson Laboratory, RRID:IMSR_JAX:005557) mice were used. CRC-001 was from Jackson Laboratory (Cat# TM00849) and XST-GI-010 was established in our laboratory at UTHealth with patient informed written consent. The studies were conducted in accordance with recognized ethical guidelines (i.e., Declaration of Helsinki, Belmont Report, and U.S. Common Rule) and appropriate approvals were obtained by the UTHealth Houston Institutional Review Board (HSC-MS-20-0327, HSC-MS-21-0074). Female 6-8-week-old mice were implanted subcutaneously with 2-3 mm PDX fragments into the right flank. When tumors reached 100–200 mm^3^, mice were randomized based on equivalent average tumor volume for each treatment group. No blinding to group allocation was performed and sample sizes were determined based on prior experience with the models to reach statistical significance. Mice were intraperitoneally dosed weekly for a total of three doses with PBS vehicle, CTX, 8E11-CPT2, R20-CPT2, or CTX in combination with 8E11-CPT2 or R20-CPT2 as indicated. Mice were routinely monitored for morbidity and mortality. Tumor volumes were measured bi-weekly and estimated by the formula: tumor volume = (length/2 × width^2^). Mice were euthanized when tumor diameter reached 15 mm. Percentage of tumor growth inhibition (% TGI) was calculated using the formula [1 – (change of tumor volume in treatment group/change of tumor volume in vehicle group)] × 100 (%).

### Statistical Analysis

All data were analyzed using GraphPad Prism 10 software (RRID:SCR_002798). Data presented as mean±SD. In vitro viability data were analyzed using the logistic nonlinear regression model. Each condition was tested in 3 independent experiments unless otherwise noted. Statistical significance of in vitro and in vivo experiments was analyzed with ANOVA and Tukey’s multiple comparison test. Kaplan-Meier analysis was performed to determine significance of treatments on survival. p < 0.05 was deemed significant.

