## Supplementary Tables S1-3 for "Cetuximab increases LGR5 expression and augments LGR5-targeting antibody-drug conjugate efficacy in patient-derived colorectal cancer models"

| **Cancer cell line/ PDX model** | **KRAS** | **NRAS** | **PIK3CA** |
| --- | --- | --- | --- |
| LoVo | G13D | WT | WT |
| AGS | G12D | WT | E453K/E545A |
| HCT116 | G13D | WT | H1047R |
| RKO | WT | WT | H1047R |
| LIM1215 | WT | WT | WT |
| DLD-1 | G13D | WT | E545K |
| SW620 | G12V | WT | WT |
| XST-GI-010 | G13D | WT | E545K |
| CRC-001 | WT | G12D | WT |

**Table S1. RAS and PIK3CA mutations in cancer cell lines and PDX models.**  Cancer cell mutations based on CCLE data and XST-GI-010 and CRC-001 PDX mutations based on next-generation sequencing analysis.

| **Average IC_50_ values (μg/ml) for 8E11-CPT2** | |
| --- | --- |
| 293T | ND |
| 293T-LGR5 | 0.035 +/- 0.023 |
| AGS | 9.97 +/- 2.27 |
| SW620 | 9.86 +/- 1.35 |
| LoVo | 12.11 +/- 5.97 |
| LIM1215 | 19.96 +/- 4.83 |
| DLD-1 | 33.57 +/-7.38 |
| HCT116 | ND |
| RKO | ND |

**Table S2. Average IC_50_ values for 8E11-CPT2 in a panel of gastrointestinal cancer cell lines.** Data representative of 2-3 independent experiments performed in triplicate. ND, not determined.

| **Loewe Synergy Scores** | | | | | | | | | | | | | | | | | | | |
| --- | --- | --- | --- | --- | --- | --- | --- | --- | --- | --- | --- | --- | --- | --- | --- | --- | --- | --- | --- |
| **Treatments** | | **LIM1215** | | | | **LoVo** | | | | | | | | | | **AGS** | | | |
| **[CTX] (μg/ml**) | | **0.001** | **0.005** | **0.01** | **0.05** | **0.1** | **0.5** | | **1** | | **2** | **5** | | **10** | **1** | | **5** | **10** | **20** |
| **[8E11-CPT2] (μg/ml)** | **60** | 2.03 | 7.58 | 12.2 | 18.47 | 29.62 | 20.41 | | 22.27 | | 1.01 | 24.33 | | 16.12 | 2.08 | | 1.48 | 0.74 | 4.13 |
|  | **20** | -3.53 | -0.47 | 9.45 | 18.44 | 26.62 | 29.95 | | 22.43 | | 8.61 | 24.74 | | 18.97 | 0.69 | | 3.58 | -0.53 | 12.45 |
|  | **6.67** | -2.25 | 2.4 | 8.02 | 15.5 | 29.95 | 21.66 | | 19.19 | | 11.25 | 21.26 | | 13.5 | 1.84 | | -0.73 | 0.53 | 13.22 |
|  | **2.22** | -1.12 | 1.7 | 7.34 | 10.19 | 32.41 | 26.61 | | 31.5 | | 10.24 | 19.91 | | 10.41 | 2.49 | | -2.51 | -4.23 | 6.21 |
|  | **0.74** | -1.35 | 0.41 | 7.82 | 1.94 | 34.07 | 29.38 | | 24.35 | | 6.03 | 16.01 | | 4.98 | 1.25 | | 1.19 | -5.47 | 0.83 |
|  | **0.25** | NT | | | | 33.76 | 27.59 | | 22.03 | | 3.05 | 12.3 | | 6.24 | -1.79 | | -1.52 | -5.13 | -2.77 |
|  | **0.082** | -0.64 | -2.17 | -4.51 | -3.15 | 4.41 | -10.3 | | 12.96 | | -3.55 | -12.95 | | -19.1 | NT | | | | |
|  | **0.027** | NT | | | | 20.71 | 4.16 | | 8.91 | | -2.01 | 6.1 | | -25.38 |  |  |  |  |  |
| **Treatments** | | **DLD-1** | | | | **SW620** | | | | | | | | | |  | | | |
| **[CTX] (μg/ml**) | | **1** | **5** | **10** | **20** | **0.1** | | **1** | | **5** | | | **10** | | |  |  |  |  |
| **[8E11-CPT2] (μg/ml)** | **60** | -3.79 | 7.17 | 7.33 | 0.9 | -2.69 | | -3.35 | | -3.18 | | | 4.23 | | |  |  |  |  |
|  | **20** | 5.34 | 35.74 | 21.52 | 23.62 | 3.58 | | -12.49 | | -8.17 | | | -7.74 | | |  |  |  |  |
|  | **6.67** | 10.72 | 11.36 | 30.46 | 11 | -13.76 | | -12.86 | | -11.48 | | | -7.68 | | |  |  |  |  |
|  | **2.22** | -1.61 | -6.44 | -2.44 | -8.68 | -3.52 | | -8.9 | | -6.49 | | | 2.65 | | |  |  |  |  |
|  | **0.74** | -3.25 | -6.95 | -12.43 | -16.68 | -8.84 | | -7.99 | | -16.18 | | | -4.96 | | |  |  |  |  |

**Table S3. Loewe synergy scores for CTX in combination with 8E11-CPT2 ADC.** NT, not tested.
