## Supplementary figures and images for "Cetuximab increases LGR5 expression and augments LGR5-targeting antibody-drug conjugate efficacy in patient-derived colorectal cancer models"

### Supplemental Figure S1

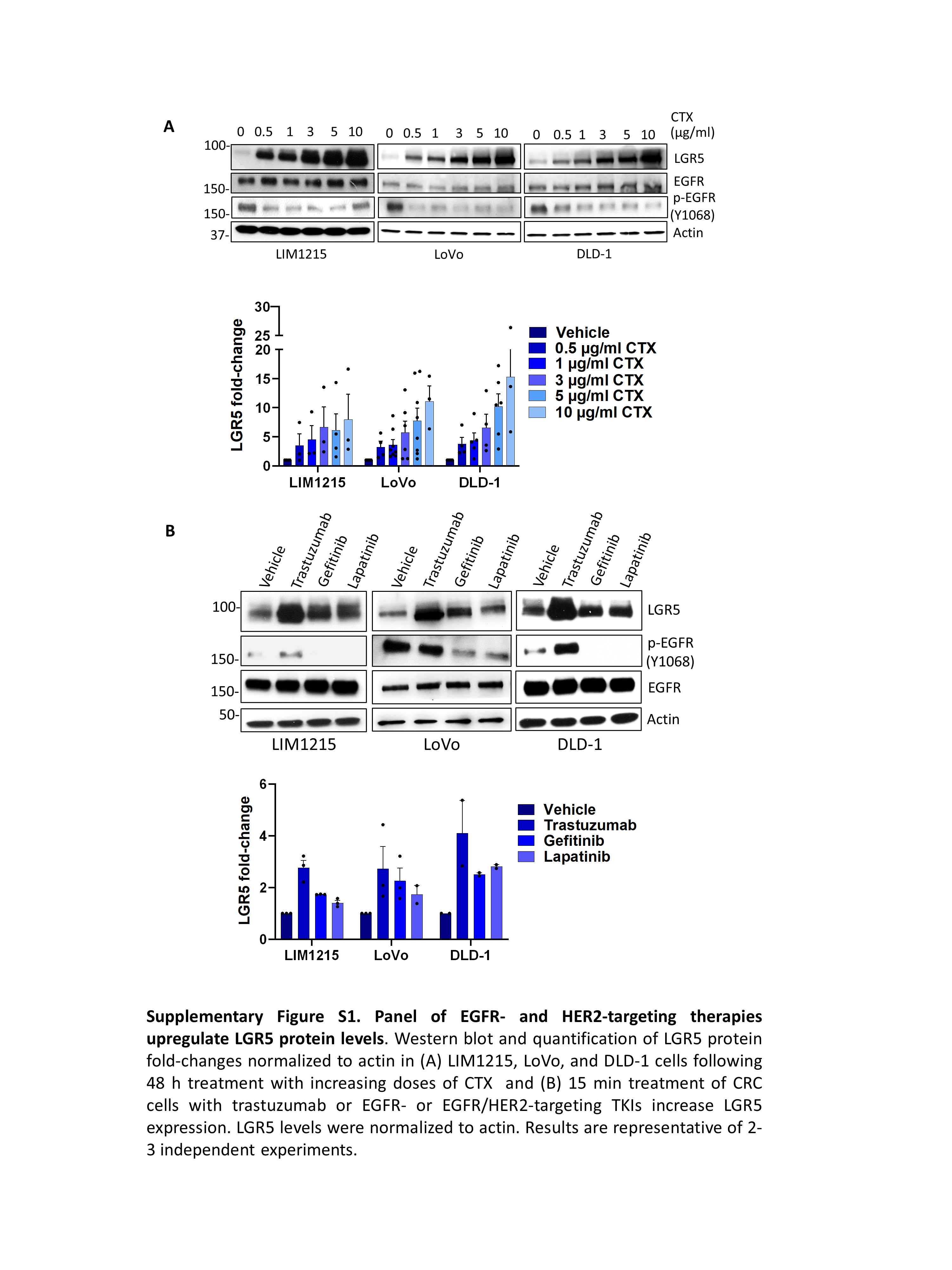

### Supplemental Figure S2

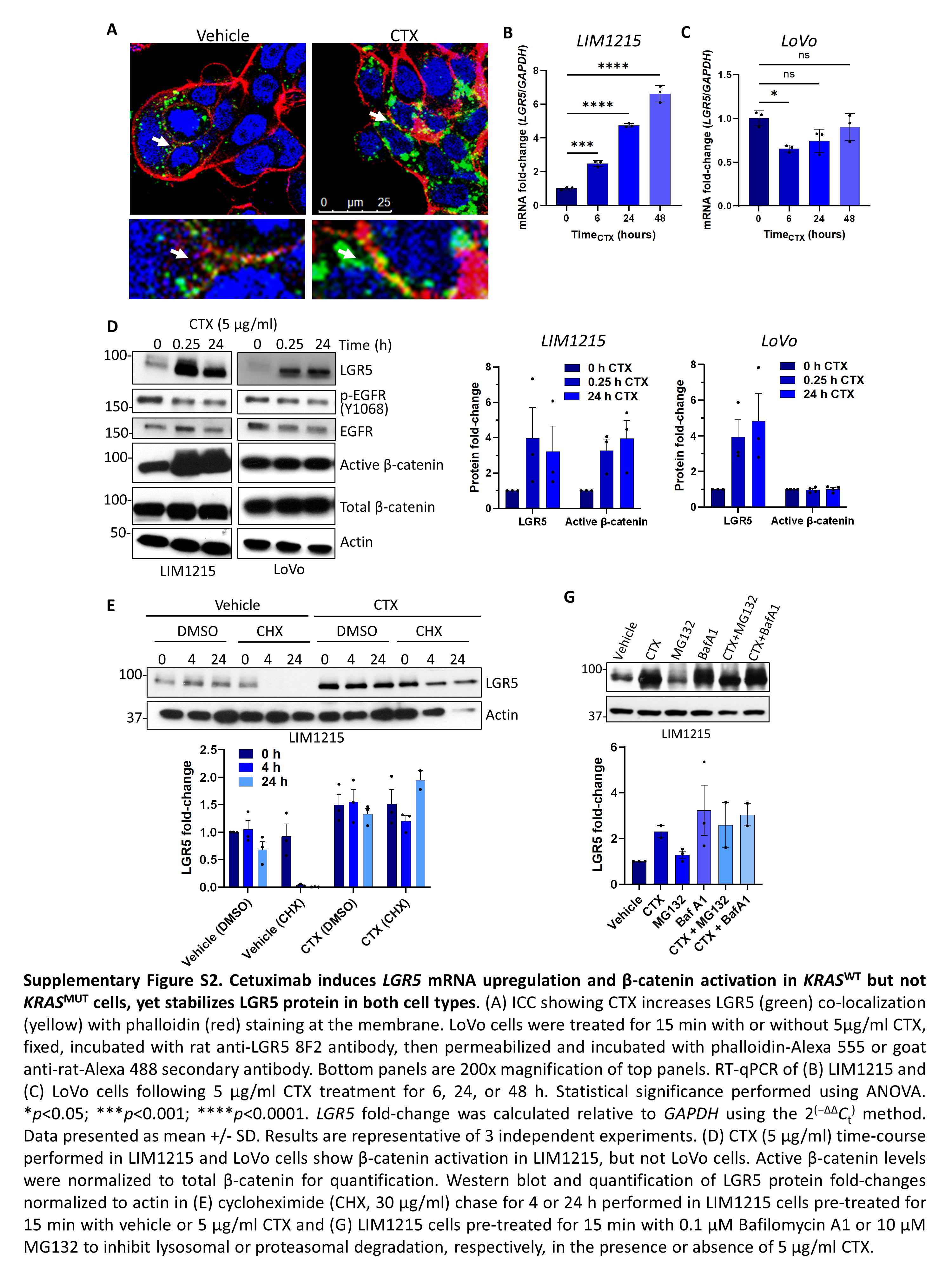

### Supplemental Figure S3

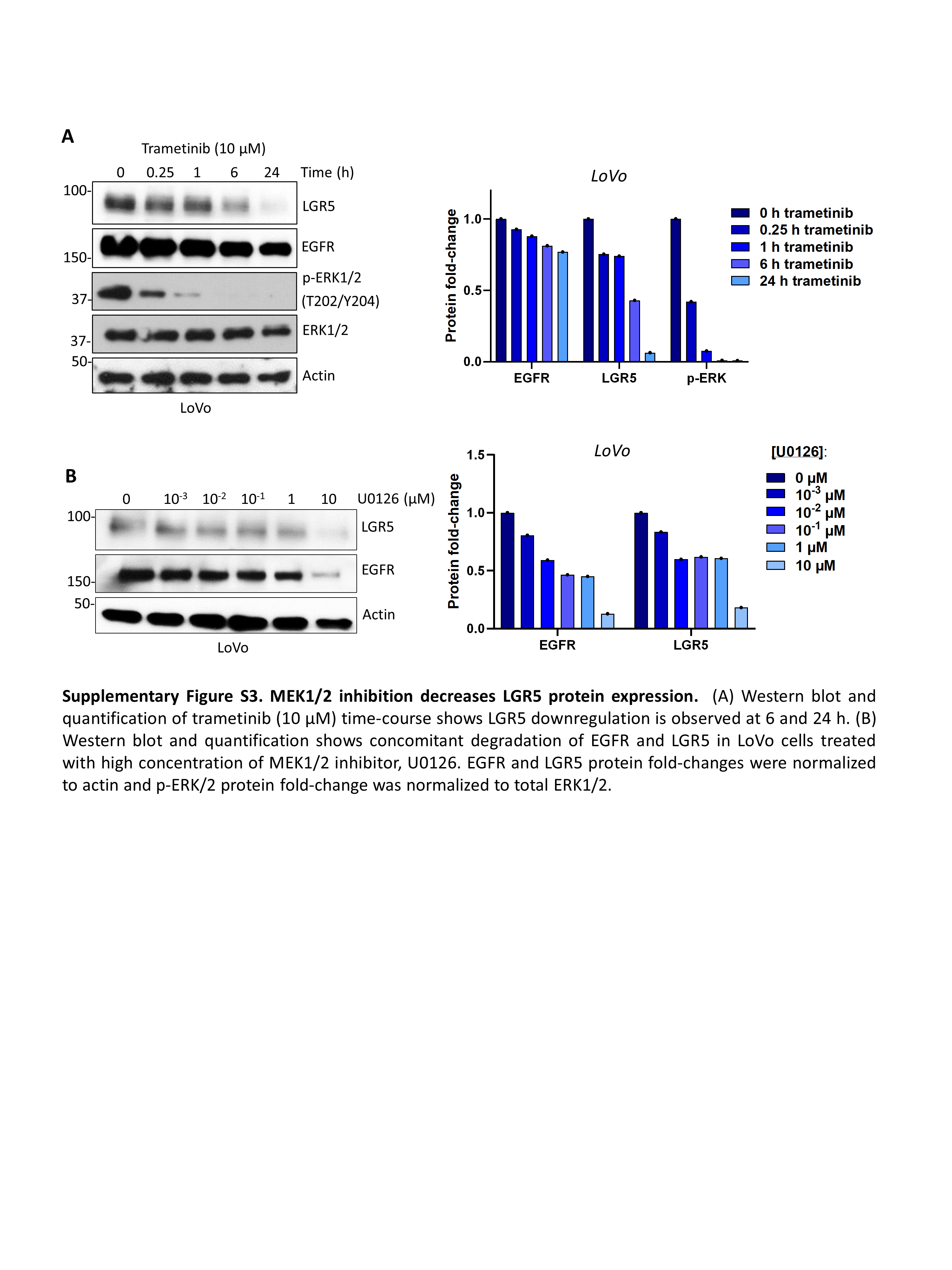

### Supplemental Figure S4

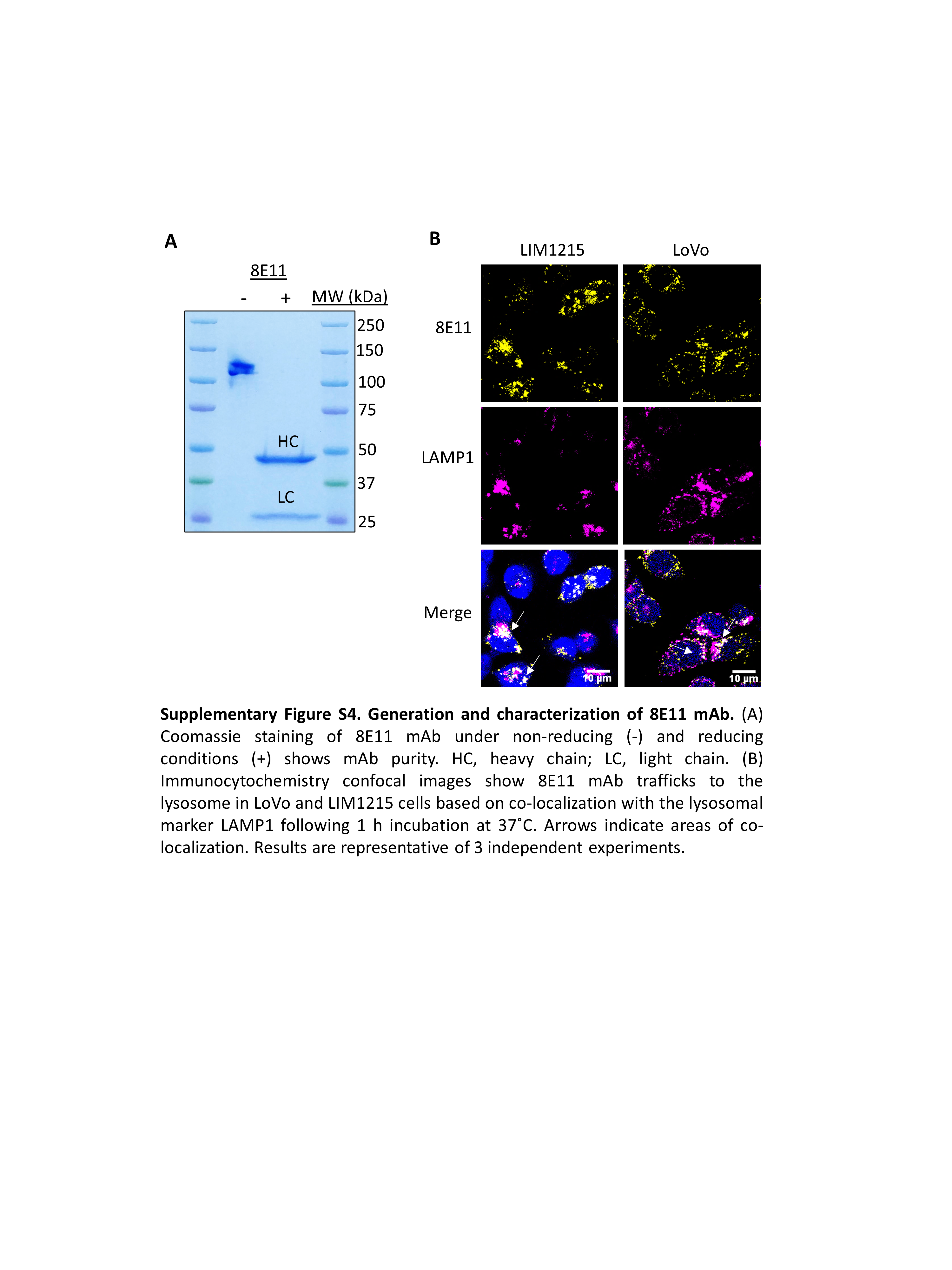

### Supplemental Figure S5

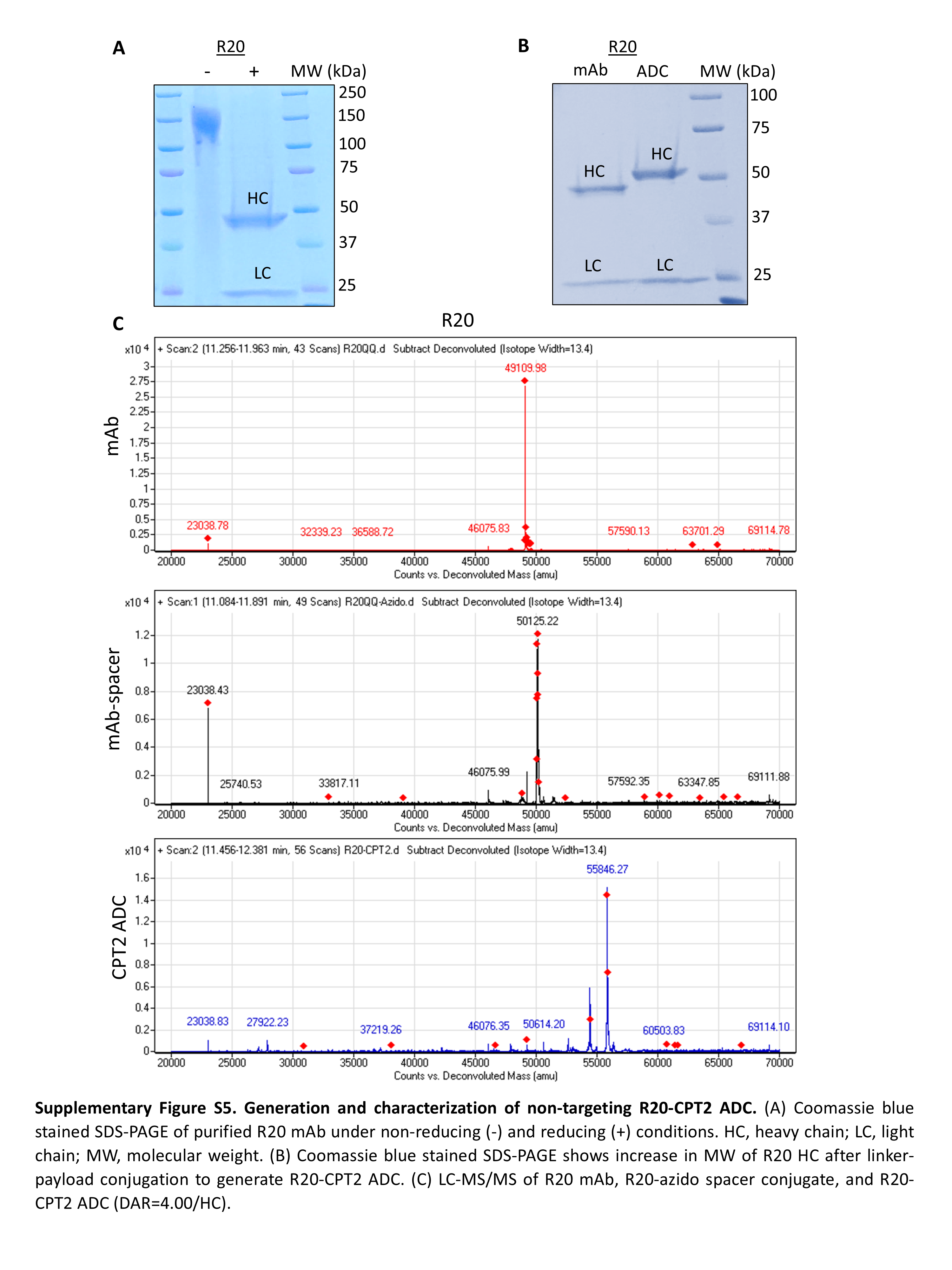

### Supplemental Figure S6

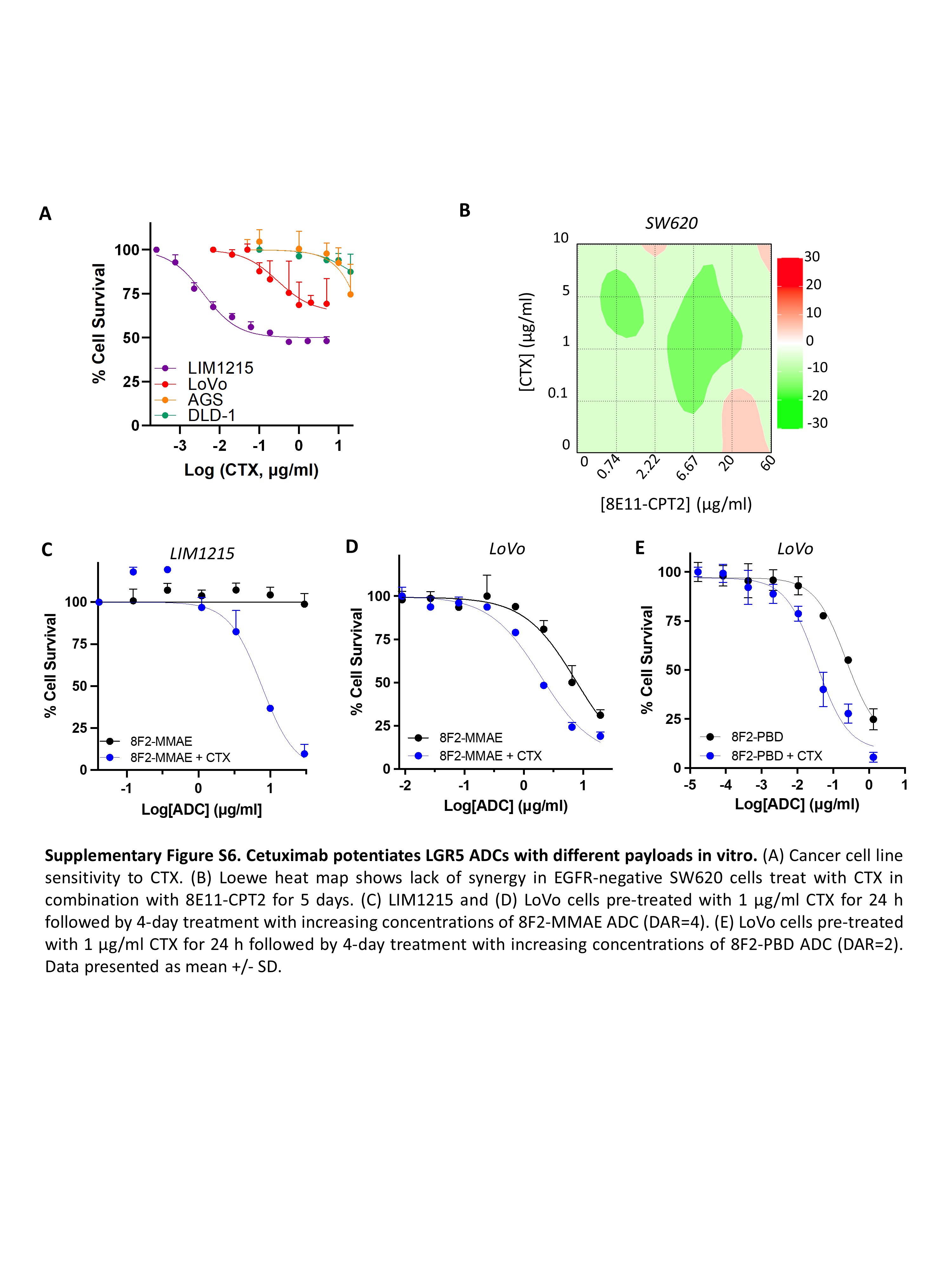

### Supplemental Figure S7

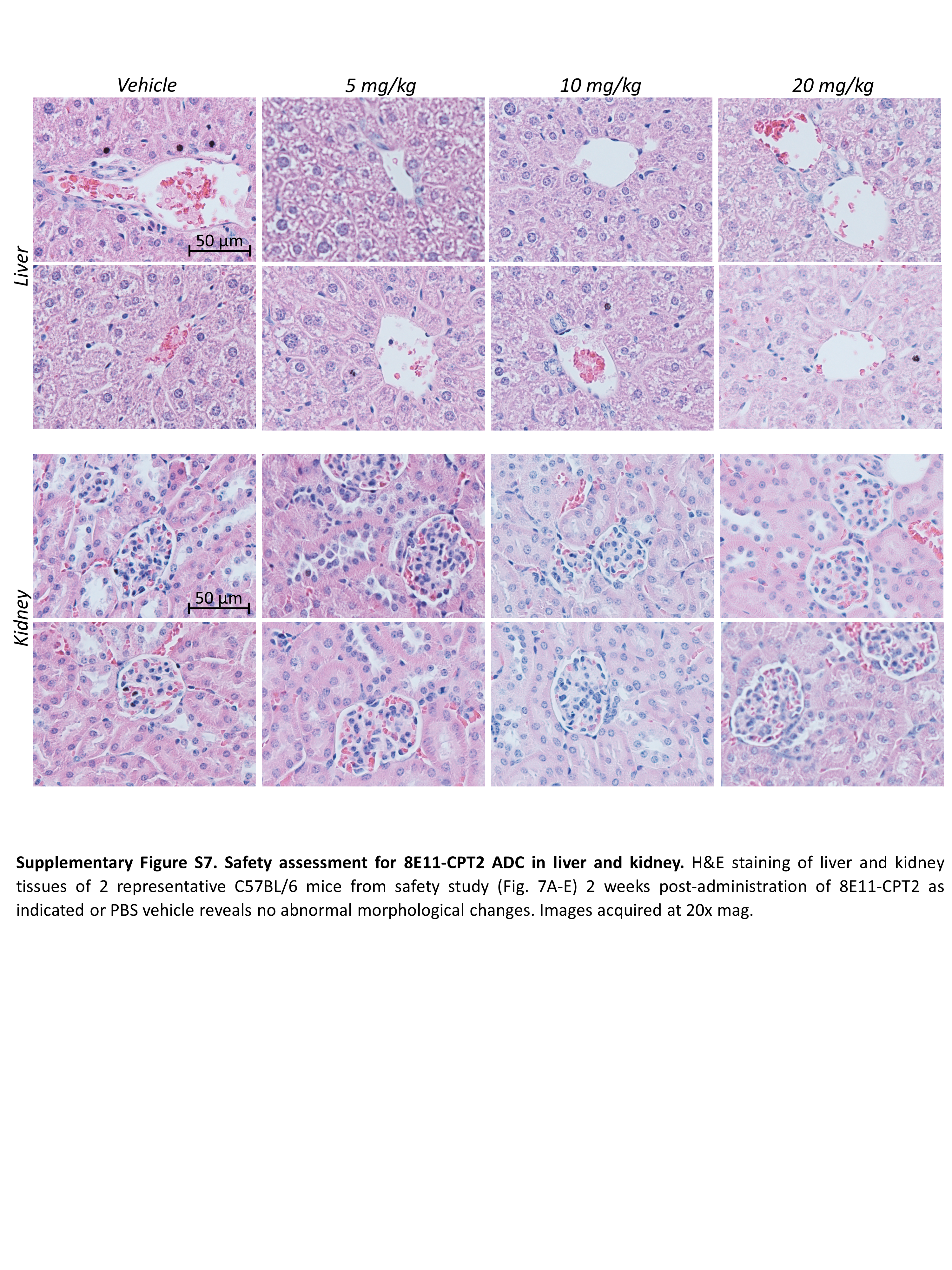

### Supplemental Figure S8

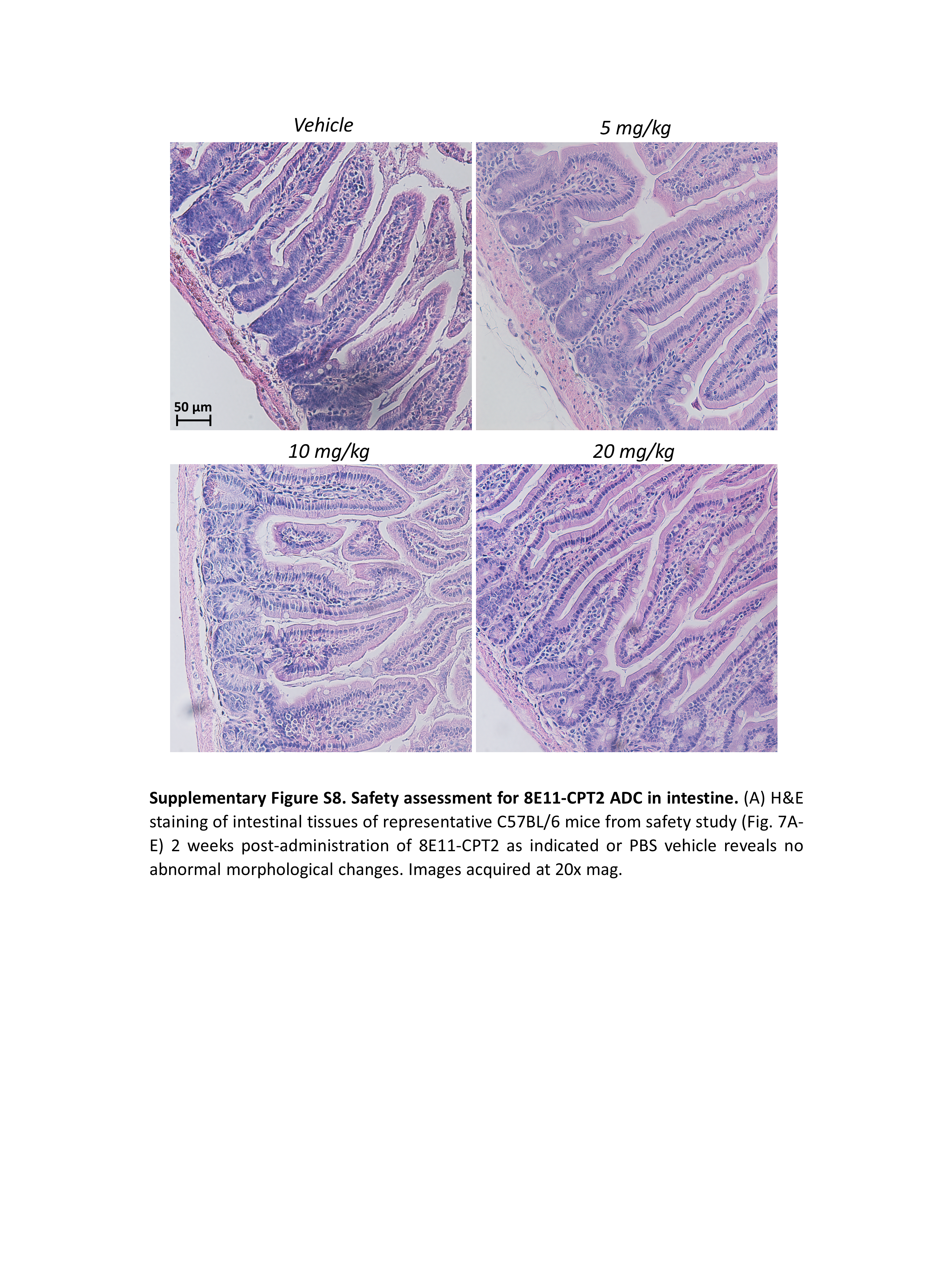

### Supplemental Figure S9

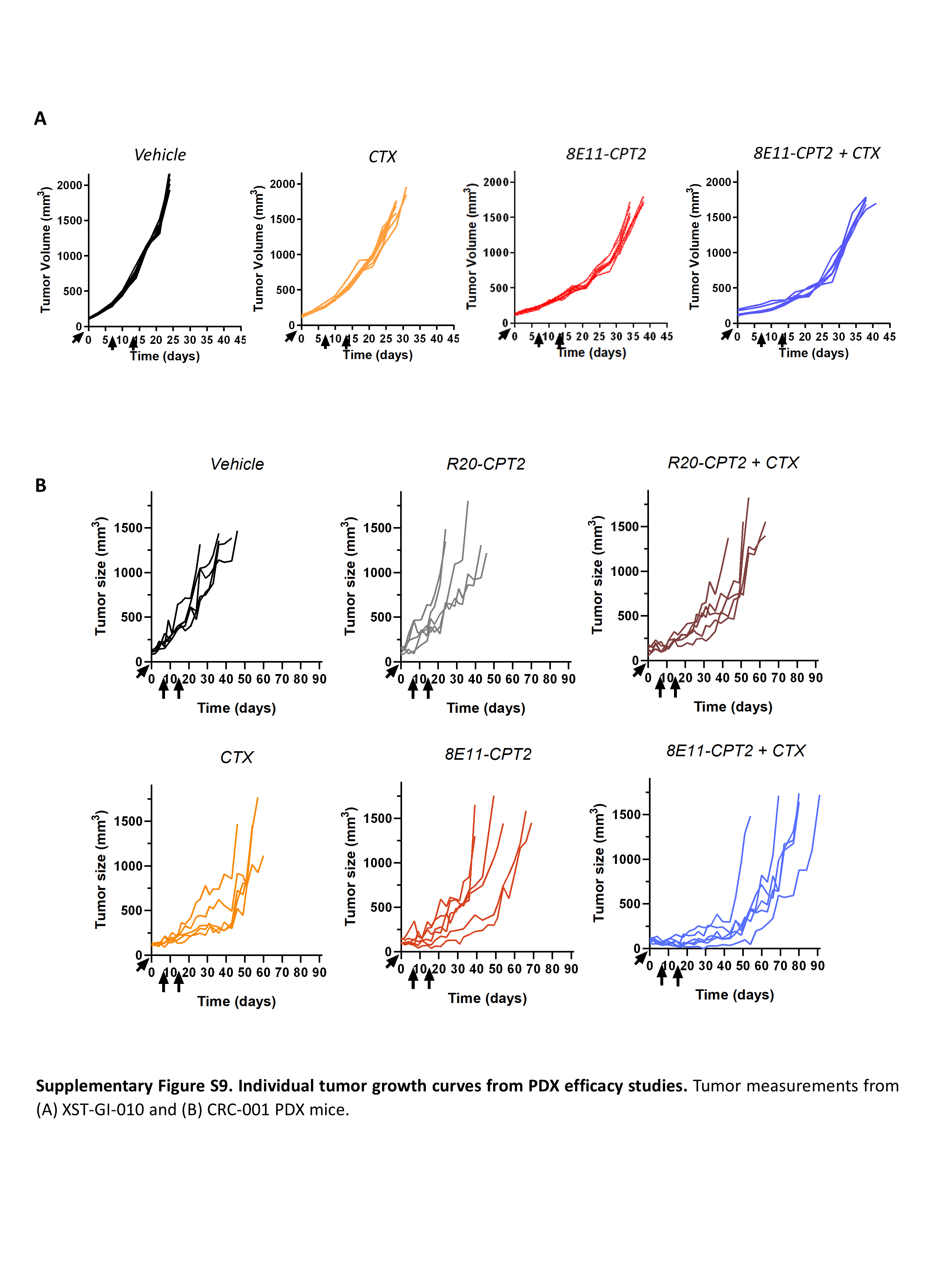

### Supplemental Figure S10

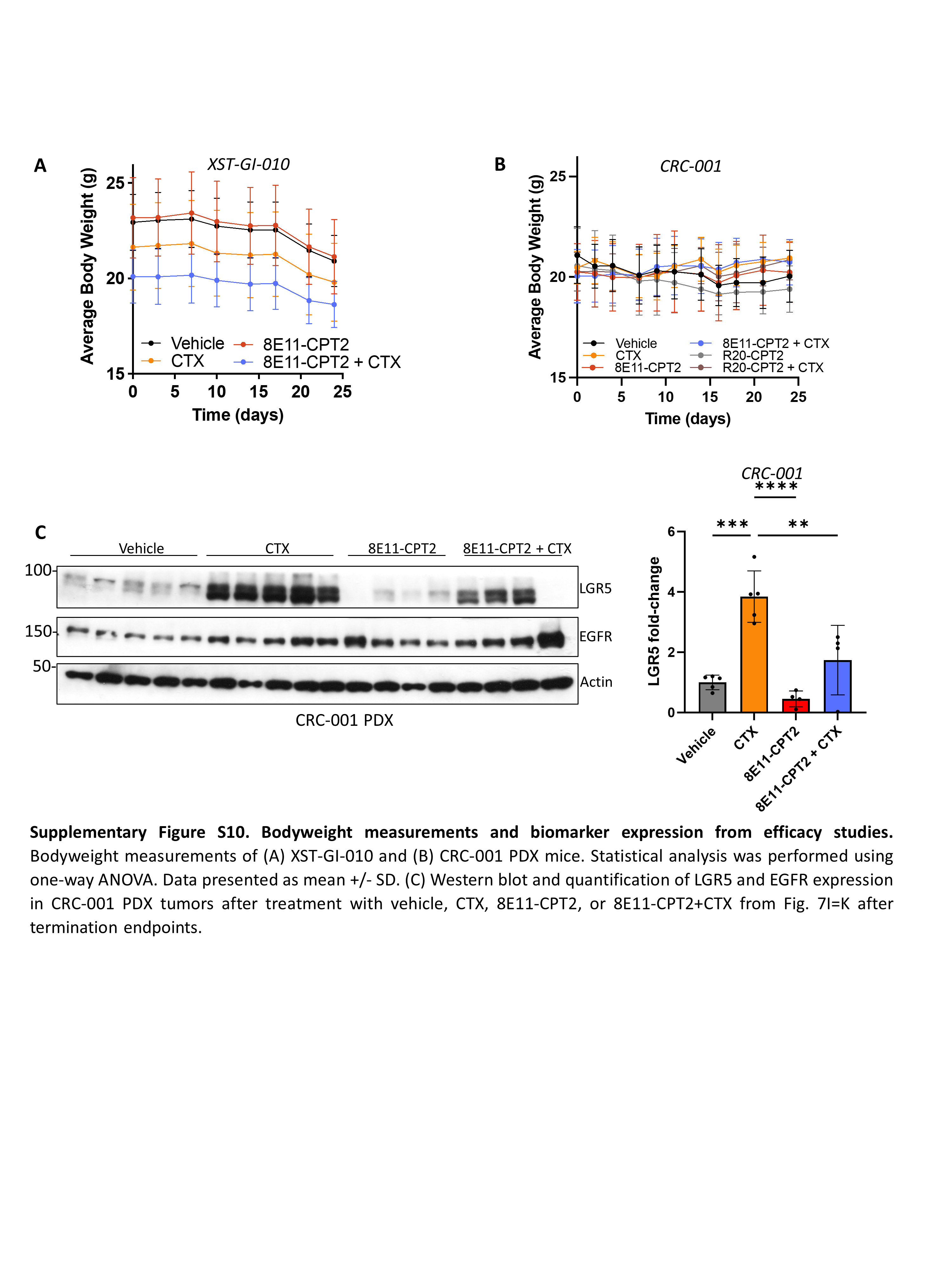
